# A harmonized public resource of deeply sequenced diverse human genomes

**DOI:** 10.1101/2023.01.23.525248

**Authors:** Zan Koenig, Mary T. Yohannes, Lethukuthula L. Nkambule, Xuefang Zhao, Julia K. Goodrich, Heesu Ally Kim, Michael W. Wilson, Grace Tiao, Stephanie P. Hao, Nareh Sahakian, Katherine R. Chao, Mark A. Walker, Yunfei Lyu, gnomAD Project Consortium, Heidi L. Rehm, Benjamin M. Neale, Michael E. Talkowski, Mark J. Daly, Harrison Brand, Konrad J. Karczewski, Elizabeth G. Atkinson, Alicia R. Martin

**Affiliations:** Stanley Center for Psychiatric Research, The Broad Institute of MIT and Harvard, Cambridge, MA 02142, USA; Analytic and Translational Genetics Unit, Massachusetts General Hospital, Boston, MA 02114, USA; Program in Medical and Population Genetics, The Broad Institute of MIT and Harvard, Cambridge, MA 02142, USA; Center for Genomic Medicine, Massachusetts General Hospital, Boston, MA 02114, USA; Department of Neurology, Massachusetts General Hospital and Harvard Medical School, Boston, MA 02114, USA; Broad Genomics, The Broad Institute of MIT and Harvard, 320 Charles Street, Cambridge, MA, 02141, USA; Data Sciences Platform, The Broad Institute of MIT and Harvard, Cambridge, MA 02142, USA; Department of Biostatistics, Harvard T.H. Chan School of Public Health, Boston, MA, USA; Institute for Molecular Medicine Finland, Helsinki, Finland; Department of Molecular and Human Genetics, Baylor College of Medicine, Houston, TX, 77030, USA

## Abstract

Underrepresented populations are often excluded from genomic studies due in part to a lack of resources supporting their analyses. The 1000 Genomes Project (1kGP) and Human Genome Diversity Project (HGDP), which have recently been sequenced to high coverage, are valuable genomic resources because of the global diversity they capture and their open data sharing policies. Here, we harmonized a high quality set of 4,094 whole genomes from HGDP and 1kGP with data from the Genome Aggregation Database (gnomAD) and identified over 153 million high-quality SNVs, indels, and SVs. We performed a detailed ancestry analysis of this cohort, characterizing population structure and patterns of admixture across populations, analyzing site frequency spectra, and measuring variant counts at global and subcontinental levels. We also demonstrate substantial added value from this dataset compared to the prior versions of the component resources, typically combined via liftover and variant intersection; for example, we catalog millions of new genetic variants, mostly rare, compared to previous releases. In addition to unrestricted individual-level public release, we provide detailed tutorials for conducting many of the most common quality control steps and analyses with these data in a scalable cloud-computing environment and publicly release this new phased joint callset for use as a haplotype resource in phasing and imputation pipelines. This jointly called reference panel will serve as a key resource to support research of diverse ancestry populations.

## Introduction

The 1000 Genomes Project (1kGP) and Human Genome Diversity Project (HGDP) have been among the most valuable genomic resources because of the breadth of global diversity they capture and their open sharing policies with consent to release unrestricted individual-level data (Bergström et al. 2020; Rosenberg et al. 2002; Li et al. 2008; 1000 Genomes Project Consortium et al. 2015, 2012). Consequently, genetic data from these resources have been routinely generated using the latest genomics technologies and serve as a ubiquitous resource of globally diverse populations for a wide range of disease, evolutionary, and technical studies. These projects are complementary; the 1000 Genomes Project is larger and has consisted of whole genome sequencing (WGS) data for many years; as such, it has been the default population genetic reference dataset, consisting of 3,202 genomes including related individuals that were recently sequenced to high coverage (Ebert et al. 2020; Byrska-Bishop et al. 2022). The 1000 Genomes Project has also been the most widely used haplotype resource, serving as a reference panel for phasing and imputation of genotype data for many genome-wide association studies (GWAS) (Lam et al. 2020; Howie et al. 2012). HGDP was founded three decades ago by population geneticists to study human genetic variation and evolution, and was designed to span a greater breadth of diversity, though with fewer individuals from each component population (Cavalli-Sforza et al. 1991; Cavalli-Sforza 2005). Originally assayed using only GWAS array data, 948 individuals have recently undergone deep WGS and fill some major geographic gaps not represented in the 1000 Genomes Project, for example in the Middle East, sub-Saharan Africa, parts of the Americas, and Oceania (Bergström et al. 2020).

The 1kGP and HGDP datasets have been invaluable separately, but far larger genomic data aggregation efforts, such as gnomAD (Karczewski et al. 2020) and TOPMed (Taliun et al. 2021), have clearly demonstrated the utility of harmonizing such datasets through the broad uptake of their publicly released summaries of large numbers of high-quality whole genomes. For example, the gnomAD browser of allele frequencies has vastly improved clinical interpretation of rare disease patients worldwide (Karczewski et al. 2017). Additionally, the TOPMed Imputation Server facilitates statistical genetic analyses of complex traits by improving phasing and imputation accuracy compared to existing resources (Taliun et al. 2021). Yet, without individual-level data access from these larger resources due to more restrictive permissions, the 1kGP and HGDP genomes remain the most uniquely valuable resources for many of the most common genetic analyses. These include genetic simulations, ancestry analysis including local ancestry inference (Maples et al. 2013), genotype refinement of low-coverage genomes (Rubinacci et al. 2021), granular allele frequency comparisons at the subcontinental level, investigations of individual-level sequencing quality metrics, and many more.

Previously, researchers wishing to combine HGDP and 1kGP into a merged dataset were left with suboptimal solutions. Specifically, the sequenced datasets had been called separately, requiring intersection of previously called sites rather than a harmonized joint-callset. Additionally, they were on different reference builds, requiring lifting over of a large dataset prior to merging, which introduces errors and inconsistencies. Here, we have created a best-in-class publicly released harmonized and jointly called resource of HGDP+1kGP on GRCh38 that will facilitate analyses of diverse cohorts. This globally-representative haplotype resource better captures the breadth of genetic variation across diverse geographical regions than previous component studies. Specifically, we aggregated these genomes into gnomAD and then jointly processed these 4,094 high-coverage whole genomes; jointly called variants consisting of single nucleotide variants (SNVs), insertions/deletions (indels), and structural variants (SVs); conducted harmonized sample, variant, and genotype quality control (QC); and separately released these individual-level genomes to facilitate a wide breadth of analyses. We quantify the number of variants identified in this new callset compared to existing releases and identify more variants as a result of joint variant calling; construct a resource of haplotypes for use as a phasing and imputation panel; examine the ancestry composition of this diverse set of populations; and publicly release these data without restriction alongside detailed tutorials illustrating how to conduct many of the most common genomic analyses.

## Results

### A harmonized resource of high-quality, high coverage diverse whole genomes

Here, we have developed a high-quality resource of diverse human genomes for full individual-level public release along with a guide for conducting the most common genetic analyses. To this end, we first extracted from gnomAD jointly called variants from 4,150 whole genomes recently sequenced to high coverage from the 1kGP and HGDP (Bergström et al. 2020; Byrska-Bishop et al. 2022), the latter of which are new to gnomAD, then harmonized project meta-data (**Table S1**). **Figure 1A** shows the locations and sample sizes of populations included in this harmonized resource. After sample, variant, and genotype QC (Chen et al. 2022), including ancestry outlier removal (**Table S2, Methods**), we identified 153,894,851 high-quality variants across 4,094 individuals, 3,400 of whom are inferred to be unrelated (**Methods, Table S3**). We computed the mean coverage within each population and project (**Figure S1-2**) as well as the mean number of SNVs per individual within each population to better understand data quality and population genetic variation (**Table S4**). While coverage was more variable among samples in HGDP (μ=34, σ=6, range=23-75X) than in 1kGP (μ=32, σ=3, range=26-66X), consistent with older samples and more variable data generation strategies (Bergström et al. 2020), all genomes had sufficient coverage to perform population genetic analysis. Consistent with human population history and as seen before (1000 Genomes Project Consortium et al. 2015), African populations had the most genetic variation with 6.1M SNVs per individual, while out-of-Africa populations had an average of 5.3M SNVs (**Table S4, Figure 1B**). The San had the most genetic variants as well as singletons per genome on average overall (**Table S4**). This is consistent with previous studies, which showed that the San had the highest genetic variation of populations studied to date, likely explained by their history traditionally as hunter-gatherers who experienced no major bottlenecks out of Africa or within Africa (Schlebusch et al. 2017; Henn et al. 2011).

**Figure 1.**
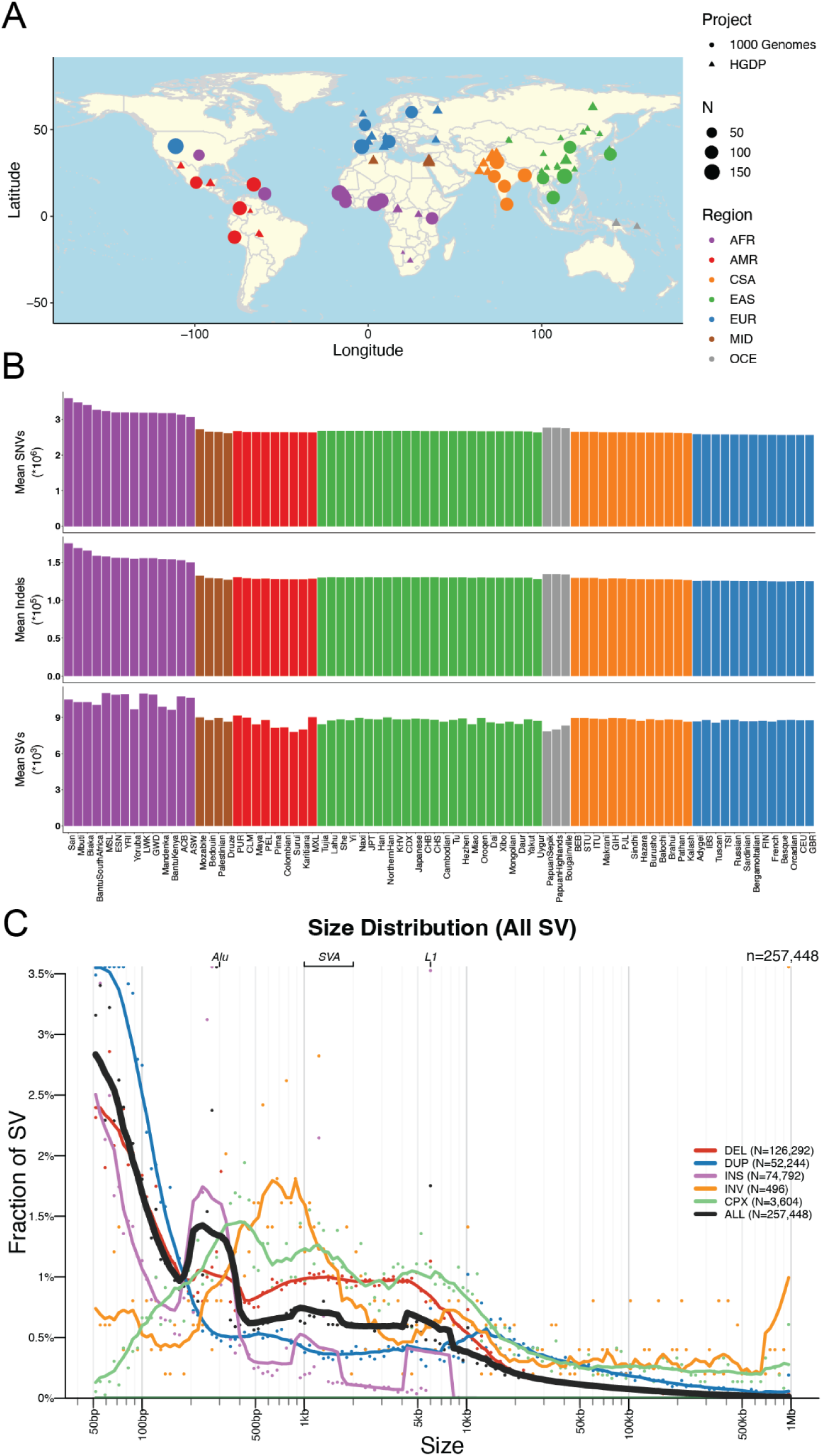
Geographical locations and genetic variants across populations. A) Global map indicating approximate geographical locations where samples were collected. Coordinates were included for each population originating from the Geography of Genetic Variants browser as well as meta-data from the HGDP (Bergström et al. 2020; Marcus and Novembre 2017). B) Mean number of SNVs (top panel), indels (2nd panel), CNVs (3rd panel), and SVs (bottom panel) per individual within each population. Colors are consistent with geographical/genetic regions in A-B), as follows: AFR=African, AMR=admixed American, CSA=Central/South Asian, EAS=East Asian, EUR=European, MID=Middle Eastern, OCE=Oceanian. C) Sizes of SVs decay in frequency with increasing size overall with notable exceptions of mobile elements, including Alu, SVA, and LINE1. Abbreviations are deletion (DEL), duplication (DUP), copy number variant (CNV), insertion (INS), inversion (INV), or complex rearrangement (CPX).

We generated a jointly genotyped structural variants (SVs) callset by detecting SVs in the HGDP genomes (**Figure S3**) using the same ensemble SV discovery tool, GATK-SV (Collins et al. 2020), as was used to generate SV calls in the high-coverage 1kGP genomes (Byrska-Bishop et al. 2022). We combined SVs from HGDP and 1kGP samples to form a non-redundant set of SV sites and uniformly genotyped them across all samples in both cohorts. In total, we identified 257,448 SV loci across 4,151 HGDP and 1kGP samples (**Figure S4A**). The frequencies of SVs were consistent with Hardy-Weinberg Equilibrium (**Figure S4F**), and distributions matched expectations from previous cohorts with the vast majority of SVs being rare (84.2% SVs at <1% allele frequency among population, **Figure S4C**). Additionally, SV size is inversely correlated with frequency (Sudmant et al. 2015; Byrska-Bishop et al. 2022; Collins et al. 2020), with notable exceptions of peaks consistent with known mobile elements, including ALU, LINE1, and SVA (**Figure 1C**). In individual genomes, we detected an average of 9,301 SVs consisting primarily of deletions (N=3,646), duplications (N=1,680), and insertions (N=3,485), as well as inversions (N=13) and complex SVs (N=109) (**Figure 1B, Figure S4B**). This data set showed comparable sensitivity to the recent 1kGP (∼9,679 SVs / genome in (Byrska-Bishop et al. 2022) and gnomAD v2 SV study (∼7,439 SVs / genome in (Collins et al. 2020)). Consistent with SNVs and indels, we observed a larger number of SVs in African populations compared to others (**Figure S5**). The precision of our SV callsets have been evaluated using 35 samples with both short-read and long-read WGS data generated by the 1kGP and the human genome structural variation consortium (HGSVC, (Ebert et al. 2021; Byrska-Bishop et al. 2022)), where 96.0% of the SVs overlapped either a short-read or long-read variant in the matched genome (**Table S6**). We observed differences in number SVs across samples from HGDP and 1kGP due to technical data generation differences, such as PCR status (**Figure S6**).

We examined global population genetic variation using principal component analysis (PCA) of the harmonized HGDP and 1kGP resource (**Figure 2**). As expected, we find PC1 differentiates AFR and non-AFR populations, PC2 differentiates EUR and EAS populations, and PC3-4 differentiate AMR and CSA populations. Subcontinental structure is also apparent in later PCs and within geographical/genetic regions (**Table S1, Figure S9-16**). These results are recapitulated with the likelihood model implemented in ADMIXTURE, where K=2 identifies similar structure in PC1, K=3 identifies similar structure in PC2, and so on (**Figure S7**). The best fit value of K=6 shown in **Figure 2** was chosen based on 5-fold cross-validation error (**Figure S8**).

**Figure 2.**
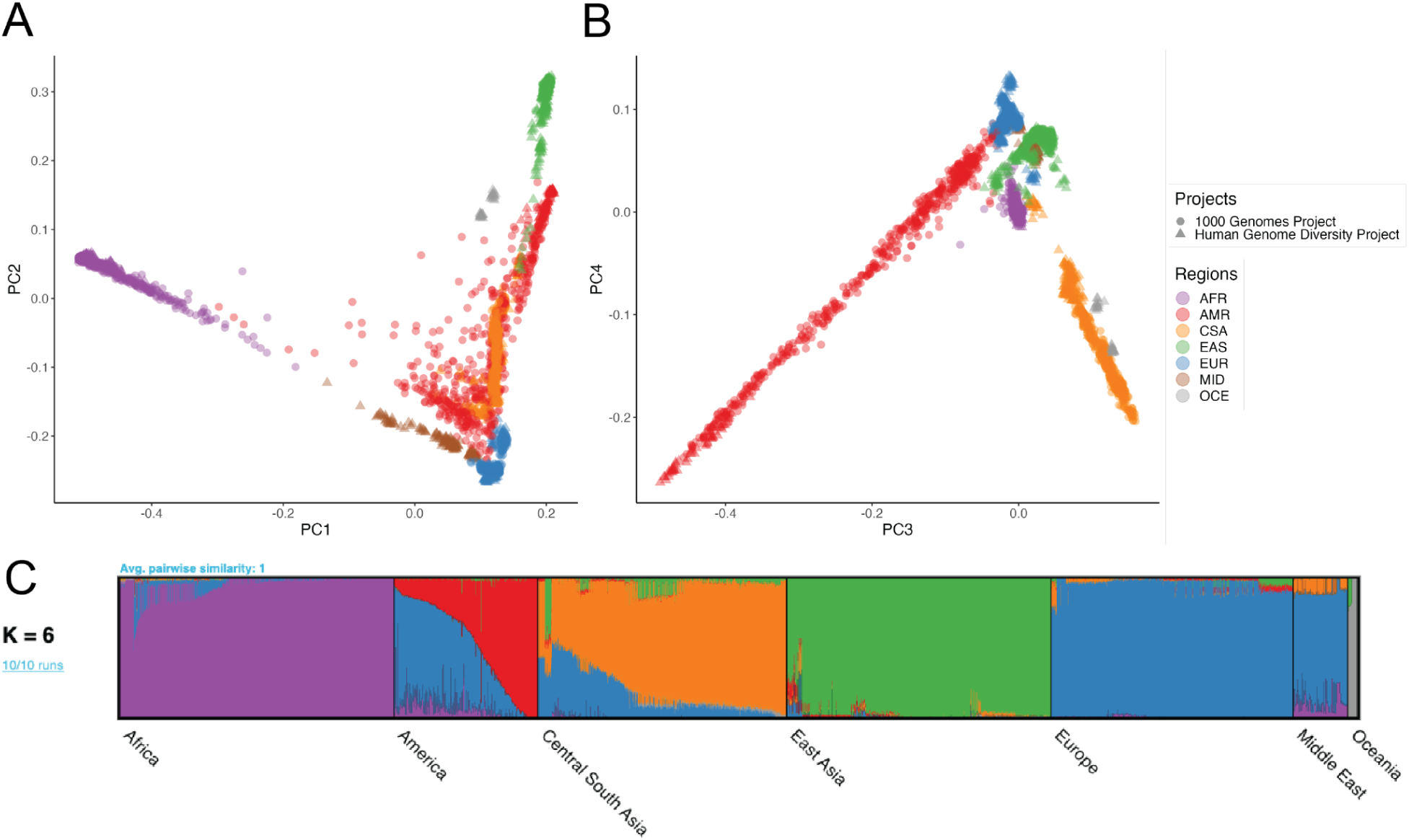
Global ancestry analysis of genetic structure in the HGDP and 1kGP resource. Regional abbreviations are as in Figure 1. A-B) Principal components analysis (PCA) plots for A) PC1 versus PC2 and B) PC3 versus PC4 showing global ancestry structure across HGDP+1kGP. Subsequent PCs separated structure within geographical/genetic regions (**Figure S9-S16**). C) ADMIXTURE analysis at the best fit value of K=6.

### Population genetic variation within and between subcontinental populations

We investigated the ancestry composition of populations within harmonized meta-data labels (AFR, AMR, CSA, EAS, EUR, MID, and OCE; **Table S1**) using PCA and ADMIXTURE analysis. Subcontinental PCA highlights finer scale structure within geographical/genetic regions (**Figure S9-S16**). For example, within AFR, the first several PCs differentiate South and Central African hunter-gatherer groups from others, then differentiate populations from East and West Africa (**Figure S10**). For AFR and AMR populations, individuals cluster similarly to the global PCA, reflecting some global admixture present in these populations (**Figure S10, S14**). The MID and OCE populations are made up of samples from the HGDP dataset only as 1kGP did not contain samples from these regions (**Figure S15-16**).

We measured population genetic differentiation using common variants with Wright’s fixation index, F_ST_ (**Figure 3A**), calculated using PLINK 1.9 (Chang et al. 2015). When populations are clustered according to pairwise F_ST_ between groups, they largely cluster by geographical/genetic region labels with a few exceptions (**Figure S18**). For example, three AMR populations are interspersed with other populations while the rest have a cluster of their own closer to the EAS populations, consistent with their population history and variable ancestry proportions that span multiple continents. Additionally, MID populations are interspersed among the EUR populations (Bedouin and Palestinian cluster together while Mozabite and Druze cluster by themselves). A CSA population, Kalash, clusters among the EUR, and an EAS population, Uygur, clusters among the CSA. AFR populations, Mbuti and San, cluster with the OCE. This mirrors their high divergence with other AFR populations and the fact that heatmap correlations are drawn from all pairwise F_ST_ estimates across populations, not just AFR. There are no interspersed populations within the other AFR cluster and no populations from AFR are interspersed among the other regions (**Table S7, Figure S18**).

**Figure 3.**
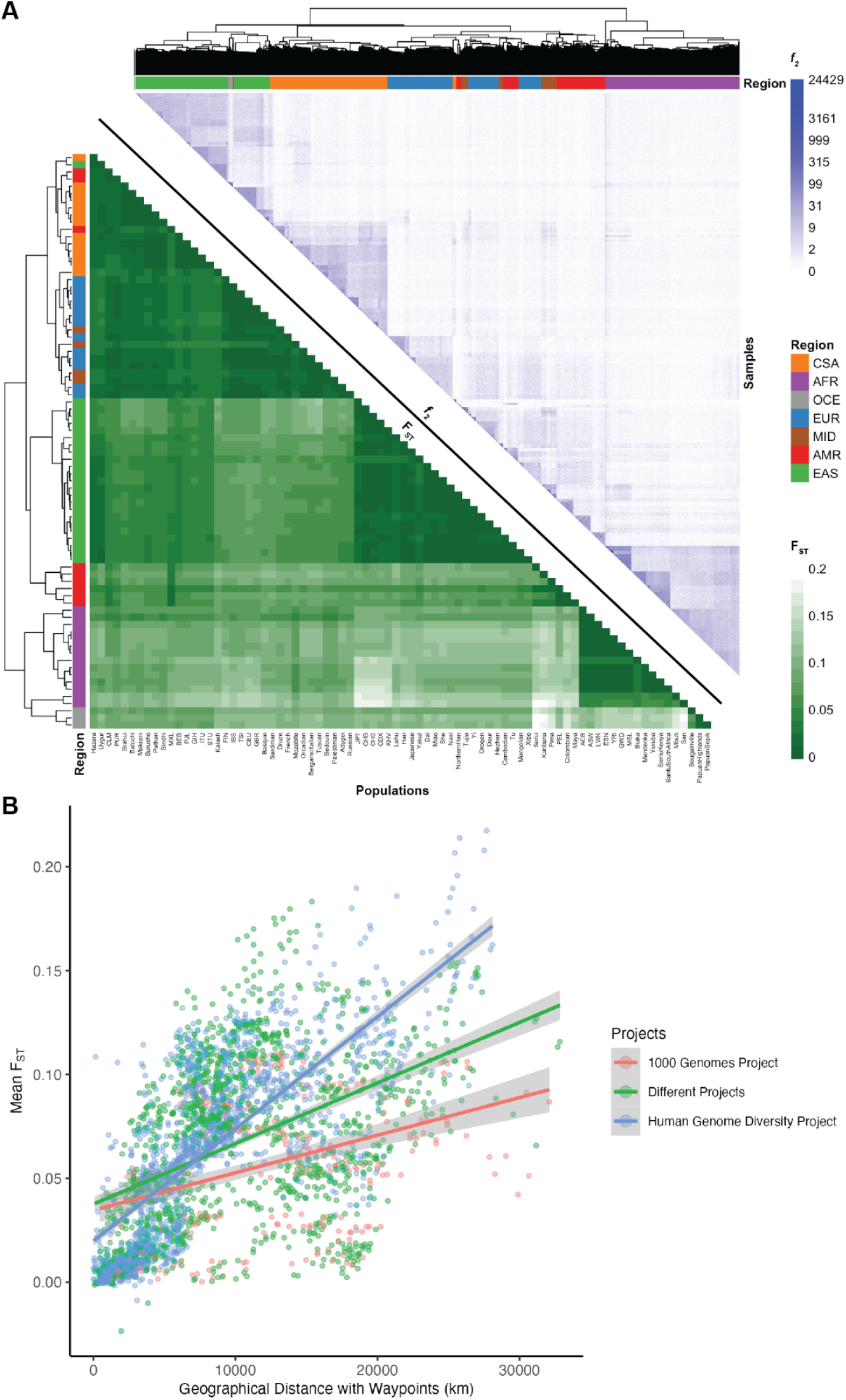
Relationships between genetic differentiation measured from common variants (F_ST_), rare variants (*f_2_*), and geography. A) Lower triangle: F_ST_ heatmap illustrating genetic divergence between pairs of populations. Upper triangle: Heatmap of *f_2_* comparisons of doubleton counts between pairs of individuals. Column and row colors at the leaves of the dendrogram show colors corresponding to meta-data geographical/genetic region and top right color bar indicates the number of doubletons shared across pairs of individuals, with more doubletons shared among individuals within the same populations and geographical/genetic regions. Interspersals of populations by meta-data labels are shown in **Figure S18** and **Table S7**. B) Genetic divergence measured by F_ST_ versus geographical distance with five waypoints calculated using haversine formula (Earth’s radius = 6,371 km).

We also compared F_ST_ versus geographical distance. We computed great circle distances using the haversine formula (Earth’s radius = 6,371 km) and pairwise geographic distances using five waypoints that reflect human migration patterns, recapitulating previous work (Ramachandran et al. 2005). The linear relationship between F_ST_ and geographical distance differs by project; specifically, HGDP has a steeper slope relating distance to F_ST_ (**Figure 3B**, likely reflecting the anthropological design intended to capture more divergent populations compared to the samples in 1kGP that reflect some of the largest populations. We compared Pearson’s correlation and Mantel tests to assess the change in the linear relationship between F_ST_ and geographical distance when incorporating waypoints. The Pearson’s correlation coefficient and Mantel statistic are both higher when waypoints are incorporated, with the highest values being when both pairs of populations are from HGDP with a correlation coefficient of 0.76 (p-value < 2.2e-16) and Mantel statistic of 0.55, p-value: 0.01 (**Table S8**).

F_ST_ measurements require group comparisons and are only based on common variants, which typically arose early in human history. Hence, we also compared rare variant sharing via pairwise doubleton counts (*f_2_*analyses, **Figure 3A**). On average, pairs of individuals within a population share 51.59 doubletons, although this varies considerably as a function of demography. For example, due to the elevated number of variants in individuals of African descent (**Figure 1**), pairs of individuals within AFR populations share on average 75.57 doubletons, whereas pairs of individuals within out-of-Africa populations share 43.8 doubletons. The individual pairs that shared the most doubletons were largely from the San population, with the top 15 sharing between 14,715 and 24,429 doubletons. Very few doubletons are shared among pairs of individuals across populations within a geographical/genetic region (u=6.79, σ=19.1) with the highest doubleton count being 4,130 between individuals from BantuSouthAfrica and San, both AFR populations. Even fewer doubletons are shared among pairs of individuals across populations from different geographical/genetic regions (u=0.79, σ=1.77) with the highest doubleton count being 638 between a pair of individuals from CDX and BEB which are EAS and CSA populations, respectively. *f_2_* clustering tends to follow project meta-data labels by geographical/genetic region, with a few exceptions.

### A catalog of known versus novel genomic variation compared to existing datasets

To demonstrate the added benefit of jointly calling these two datasets compared to using each component dataset alone, we have compiled metrics that compare our harmonized dataset with each individual dataset comprising it (Bergström et al. 2020; Byrska-Bishop et al. 2022), the previous phase 3 1kGP dataset sequenced to lower coverage (1000 Genomes Project Consortium et al. 2015), and the widely used gnomAD dataset (Chen et al. 2022). This jointly called HGDP+1kGP dataset contains 153,894,851 SNVs and indels that passed QC, whereas phase 3 1kGP has 73,257,633. High-coverage WGS of 1kGP (referred to here as NYGC 1kGP based on where they were sequenced) has 119,895,186, high-coverage WGS of HGDP (referred to here as Bergstrom HGDP based on the publication) has 75,310,370, and gnomAD has 644,267,978 high-quality SNVs and indels (Chen et al. 2022) (**Table S10**). Because gnomAD now contains both HGDP and 1kGP, we built a synthetic subset of gnomAD that removes allele counts contributed by HGDP and 1kGP. When comparing the HGDP+1kGP dataset to this synthetic version of gnomAD that excludes HGDP+1kGP, we show that variants unique to gnomAD are disproportionately rare (**Figure 4**, **Table S12**). In contrast, compared to the comprising datasets of HGDP only, NYGC 1kGP only, and phase 3 1kGP, the HGDP+1kGP dataset contributes a sizable fraction and number of variants spanning the full allele frequency spectrum, including both rare and common variants (**Figure 4**). Specifically, there are 84M novel variants (53%) in HGDP+1kG compared to HGDP only, 43M (27%) compared to NYGC 1kGP only, and 83M (53%) compared to phase 3 1kGP (**Table S12**). However, rare variants are particularly enriched; in all of the comparison datasets aside from gnomAD, the HGDP+1kGP dataset contains the largest proportion of rare variants. Few variants in the phase3 1kGP dataset were not in the HGDP+1kGP dataset or NYGC 1kGP because samples are entirely overlapping, as reported previously (Byrska-Bishop et al. 2022).

**Figure 4.**
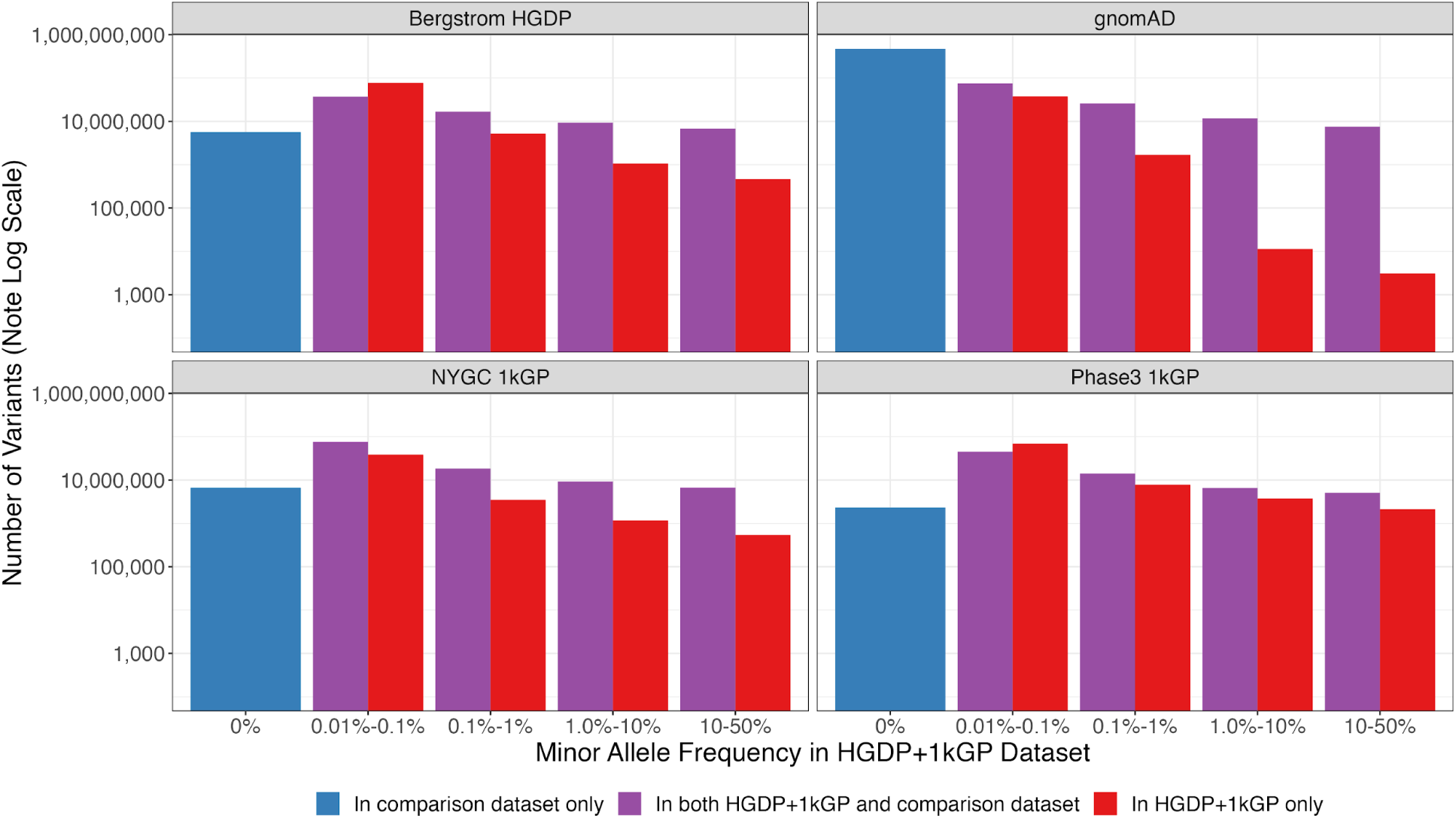
Number of variants identified in this dataset compared to commonly used existing datasets as a function of allele frequency. The number of variants on a log scale is plotted by minor allele frequency bin within the harmonized HGDP+1kGP dataset. We show variants found in the harmonized HGDP+1kGP dataset only (red), variants shared between the harmonized dataset and each comparison dataset (purple), and variants that are only found in each comparison dataset (blue). More information on exact numbers and comparisons by QC within and across datasets can be found in **Supplementary Table S11-12**.

### Phased haplotypes improve phasing and imputation accuracy and flexibility compared to existing public resources

We next developed the HGDP+1kGP dataset as a haplotype resource by phasing variants together using SHAPEIT5 (Hofmeister et al. 2023), including information about trios (**Methods**). We first evaluated phasing switch error rate using 34 genomes that overlapped with 1kGP and had fully phased genome assemblies including long read sequencing data in the Human Genome Structural Variation Consortium, Phase 2 (HGSVC2) (Ebert et al. 2021). We treated the HGSVC2 genomes as “truth”, then compared statistical phasing with the HGDP+1kGP versus 1kGP reference panel. We find lower switch error rate when using HGDP+1kGP versus 1kGP only in both SNPs (mean=0.00184, sd=0.00145 in HGDP+1kGP vs mean=0.00338, sd=0.00327 in 1kGP) and indels (mean=0.00899, sd=0.00627 in HGDP+1kGP vs mean=0.0148, sd=0.0112 in 1kGP, **Figure 5A**).

**Figure 5.**
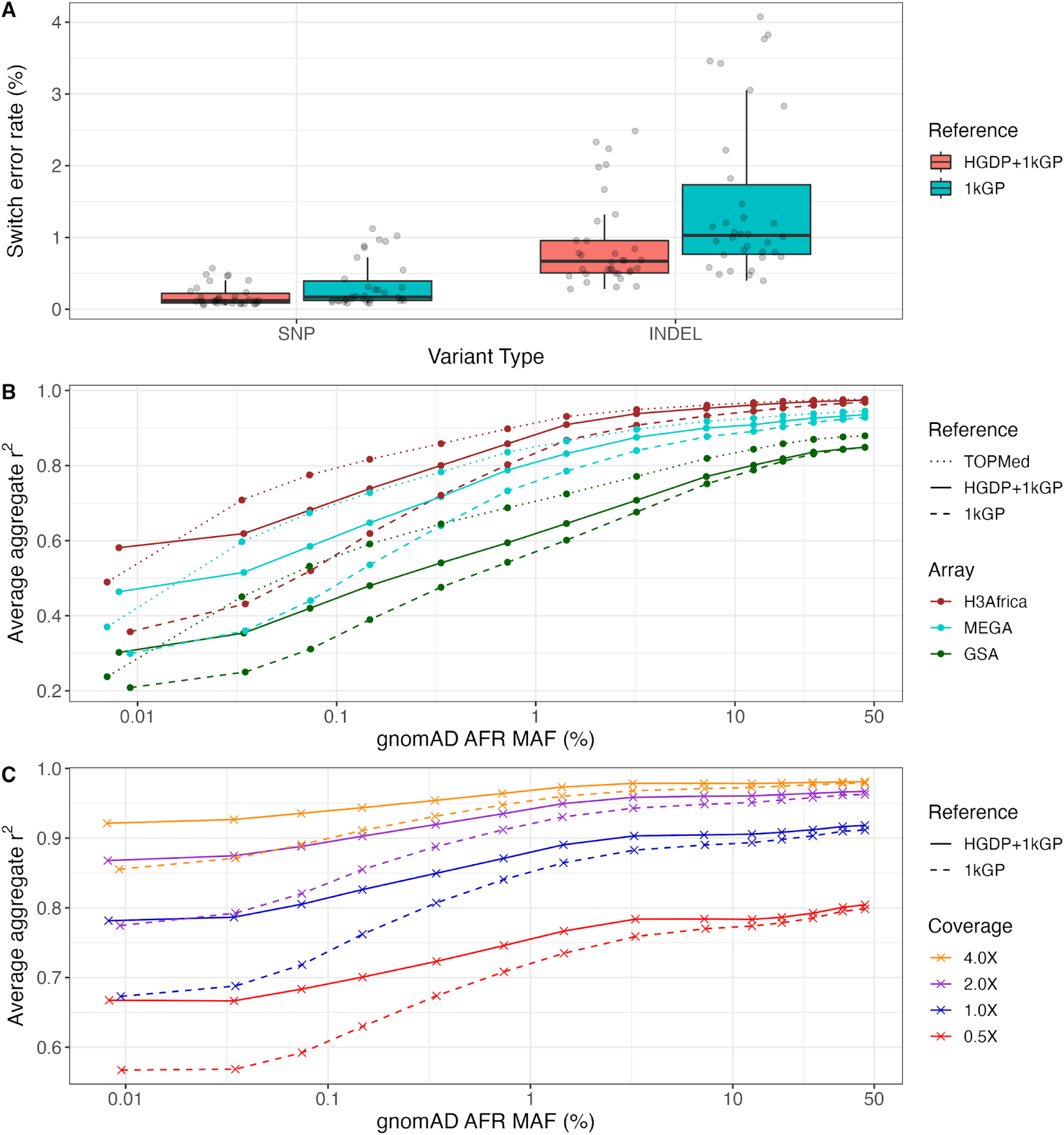
Phasing and imputation accuracy are improved across data generation strategies compared to existing reference panels. A) Switch error rates for SNPs and indels in a truth set of 34 HGSVC2 genomes when using HGDP+1kGP versus 1kGP reference panels for phasing. B-C) Imputation performance as a function of minor allele frequency (MAF) for AFR in gnomAD v3.1 data using TOPMed, HGDP+1kGP, and 1kGP reference panels in B) SNP array and C) low-coverage sequencing data. Aggregate r^2^, which is the correlation between the imputed dosages and high coverage “truth” genotype calls, was computed in MAF bins and averaged across chromosomes 1-22. The validation set is composed of 93 AFR individuals sequenced at 30X coverage (Martin et al. 2021).

We next evaluated imputation accuracy with common genetic data generation strategies. We used 93 downsampled whole genomes sequenced as part of the NeuroGAP Project consisting of East and South African participants to either 1) sites on relevant and commonly used GWAS arrays (Illumina GSA, MEGA, and H3Africa), or 2) lower depths of coverage (0.5X, 1X, 2X, and 4X), as done previously (Martin et al. 2021). We imputed GWAS arrays using IMPUTE5 (Rubinacci et al. 2020) and low-coverage genomes using GLIMPSE (Rubinacci et al. 2021). With several haplotype reference panels, we compared average imputation accuracy as a function of allele frequency estimated from gnomAD AFR frequency given our small sample size. For arrays, we compared imputation accuracy using NYGC 1kGP, HGDP+1kGP, and TOPMed imputation panels (Kowalski et al. 2019). As expected with GWAS arrays, HGDP+1kGP improves accuracy compared to 1kGP but not compared to the much larger TOPMed data (**Figure 5B**). Because low-coverage genomes require individual-level haplotypes for imputation that ideally operate on genotype likelihoods rather than initial genotype calls (Rubinacci et al. 2021), we were unable to compare the TOPMed panel. For low-coverage genomes, we find much higher imputation accuracies with HGDP+1kGP. At rarer variants, the imputation accuracy differences due to reference panel used are almost as high as those due to higher depths of sequencing; for example, rare variants sequenced to 2X depth and imputed with HGDP+1kGP are imputed almost as accurately as rare variants sequenced to 4X depth and imputed with 1kGP, thus highlighting the utility of the larger sample size and diversity in this resource (**Figure 5C**).

### Facilitating broad uptake of HGDP+1kGP as a public resource via development of detailed tutorials

In an effort to increase accessibility of this dataset, we have made publicly available tutorials of our analyses implemented primarily in Hail (https://hail.is). Hail is an open source, Python-based, scalable tool for genomics that enables large-scale genetic analyses on the cloud. Tutorials can be accessed through Github via iPython notebooks (https://github.com/atgu/hgdp_tgp/tree/master/tutorials), and all underlying datasets are publicly available in requester-pays Google Cloud Platform buckets.

These tutorials cover various aspects of QC and analysis, including sample, variant, and genotype QC; visualizing distributions of QC statistics by meta-data labels across diverse populations; filtering variants using LD, allele frequency, and missingness information; inferring relatedness; running PCA to infer ancestry; computing descriptive statistics including variant counts and coverage metrics; conducting population genetic analyses; and intersecting external datasets with HGDP+1kGP as a reference panel to apply ancestry models and infer meta-data labels (**Figure 6**). For example, we intersected the publicly available Gambian Genome Variation (GGV) Project sequenced to low coverage with the HGDP+1kGP resource, trained a random forest on HGDP+1kGP geographical/genetic region meta-data labels, then applied this model to the GGV data to determine ancestry labels, which were all inferred to be AFR (**Figure S17**). When intersecting external datasets to apply ancestry labels, an important consideration is how many variants must overlap and how much missingness is tolerated to project external samples into the same PCA space as the reference panel and assign meta-data labels given PCA shrinkage (Dey and Lee 2019). We find that <5% missingness is typically required to accurately assign ancestry labels (**Figure S20, Table S13**). In addition to all these analyses, we anticipate that there will be additional uses of this resource not documented in these tutorials, such as for phasing and imputation. To facilitate these uses, we have phased the HGDP+1kGP dataset and released phased haplotypes that others can use to support phasing and imputation in their own datasets. We have also developed computational pipelines implemented in GWASpy that use these phased reference haplotypes, and tested them by applying phasing and imputation to diverse samples genotyped as part of other ongoing work.

**Figure 6.**
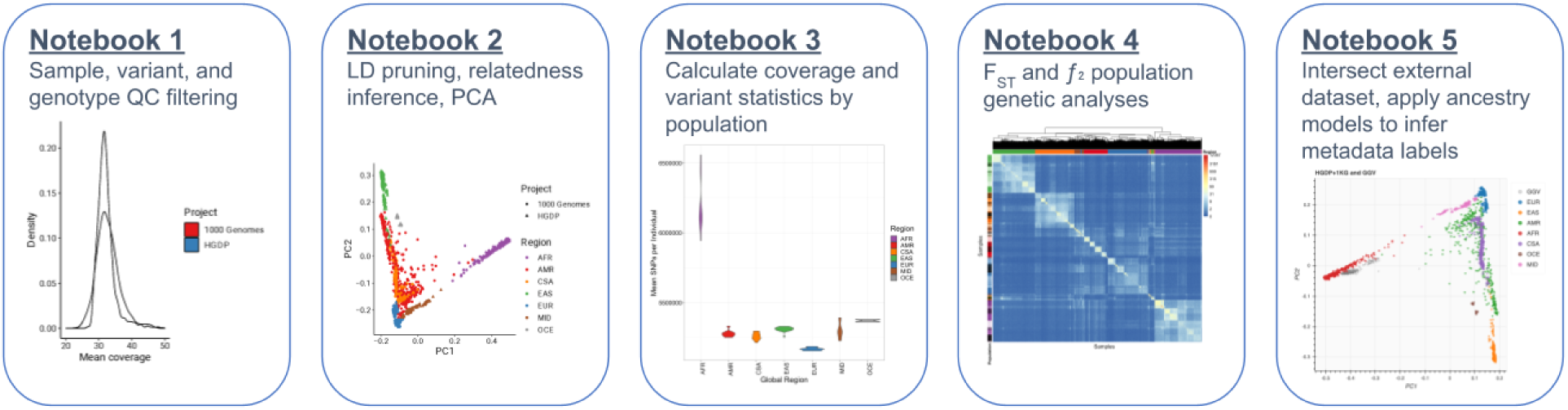
Overview of tutorials that use cloud computing to conduct common genetic data analyses. We have developed five iPython notebooks with tutorials for conducting many of the most common genetic analyses, including QC of sequencing data, relatedness inference and PCA, calculating statistics by population, analyzing genetic divergence, and applying ancestry analysis to a new dataset using HGDP+1kGP as a reference panel.

## Discussion

The 1000 Genomes Project and Human Genome Diversity Project were landmark efforts to increase the unrestricted public availability of genomic data from a geographically and ancestrally diverse set of individuals. These resources have been widely used across research efforts for decades, including as reference panels for ancestry inference, phasing, imputation, genotype refinement, and investigations into population history and demography. However, these datasets have historically been discrete, leading to suboptimal intersections when a combined analysis of all samples is required.

The harmonized variant processing, quality control, and improved coverage of variants across the allele frequency spectrum in this jointly called resource will facilitate the improved study of diverse populations. The callset formally released here has already been used as a resource of global diversity in the Genome Aggregation Database (gnomAD) (Chen et al. 2022), the Pan-UK Biobank Project (Karczewski et al.), the Global Biobank Meta-analysis Initiative (GBMI) (Zhou et al. 2022), and the Covid-19 Host Genetics Initiative (The COVID-19 Host Genetics Initiative and Ganna 2021). A primary use of this data is as a global reference for PCA – SNV loadings are freely shared so that user cohorts can be aligned to the same PC space as this reference panel. In GBMI, harmonizing ancestry analysis with this resource served as a quality control measure to ensure that ancestral groupings were applied consistently and that control for population stratification was performed adequately (Zhou et al. 2022). Building on this approach and given the critical need for greater diversity in genomic studies, sequencing centers can use this resource to build dashboards that continuously monitor the diversity of samples being sequenced in real time.

This callset is also phased for use as a haplotype resource, potentially providing higher phasing and imputation accuracy particularly for underrepresented populations. While resources such as the Haplotype Reference Consortium (HRC) and TOPMed Imputation Panel are already useful (McCarthy et al. 2016; Kowalski et al. 2019), they either provide individual-level data but lack diversity (HRC) or are very large with significant diversity but do not share individual-level data (TOPMed). This limits the application of new methods, such as those needed to support low-coverage sequencing, which is receiving growing interest as it is comparable in cost to many genotype arrays and is especially beneficial to underrepresented populations (Martin et al. 2021). Combinations of high-coverage exome and low-coverage genome sequencing are also of growing interest and could be uniquely supported by this resource. It is also being used for developing computational and analytical tools for genotype refinement and imputation, conducting data QC across varying depths of coverage, and evaluating technical biases. For example, we observed fewer SVs in the HGDP genomes than 1kGP genomes among similar ancestry groups, which was primarily explained by PCR+ and PCR-free sequencing libraries.

While this resource is more globally representative than most public datasets, certain geographic areas and ancestries are still underrepresented. HGDP, designed over two decades ago alongside the Human Genome Project, was one of the earliest studies of its kind and therefore faced some ethical controversies that remain relevant today (Resnik 1999; Greely 1999). For example, challenging issues of individual versus collective consent particularly among Amerindigenous communities parallel those currently being navigated by the *All of Us* Research Program; while criticisms have been raised, consensus has not been reached. HGDP responded to similar criticisms at the time, and developed the Model Ethical Protocol whose principles still guide all major genetic research projects to date (Weiss et al. 1997). The risk and beneficence of ongoing massive-scale efforts such as *All of Us*, whose mission is “to accelerate health research and medical breakthroughs, enable individualized prevention, treatment, and care for all of us” must be wrestled with to minimize risks and ensure adequate representativeness such that all can benefit from genomics research and ultimately precision medicine.

As genetically diverse datasets continue to grow to massive scales, it will be invaluable for researchers to be equipped with tools and resources that facilitate scalable, efficient, and equitable analysis, including allele frequency information which informs variant deleteriousness (Karczewski et al. 2017). In the service of this goal, we concurrently release a series of detailed tutorials designed to be easily accessible in iPython notebooks demonstrating many common genomic analytic techniques as implemented in the cloud-native Hail software framework, which allows for flexible, computationally efficient, and parallelized analysis of big data. The release of this resource on GRCh38 along with these detailed tutorials reduce barriers acknowledged by clinical labs which have not yet migrated to the latest genome build, citing that they do not feel the benefits outweigh the time and monetary costs and/or lack sufficient personnel to shift (Lansdon et al. 2021). The tutorials also provide a code bank for researchers to conduct a variety of analyses, including conducting quality control of whole genome sequencing data, calculating variant and sample statistics within groups, analyzing population genetic variation, and applying ancestry labels from a reference panel to their own data. Overall, resources like this are essential for empowering genetic studies in diverse populations.

## Methods

### Genetic datasets

#### Human Genome Diversity Project (HGDP)

HGDP genomes sequenced and described previously (Bergström et al. 2020) were downloaded from http://ftp.1000genomes.ebi.ac.uk/vol1/ftp/data_collections/HGDP/. Because the publicly available gVCFs were not the output of GATK HaplotypeCaller and were incompatible with joint calling, we reprocessed these genomes and conducted joint variant calling as part of gnomAD v3 (Chen et al. 2022). Most HGDP genomes were PCR-free (N=760), but some included PCR prior to sequencing (N=169). They were also sequenced at different times, for example as part of the Simons Genome Diversity Project (SGDP, N=120) or later at the Sanger Institute (N=801). More details are available from the source studies (Bergström et al. 2020; Mallick et al. 2016).

#### 1000 Genomes Project (1kGP)

1kGP genomes have been sequenced as part of multiple efforts, first to mid-coverage as phase 3 of the 1kGP (1000 Genomes Project Consortium et al. 2015) and more recently to high-coverage (≥30X) at the New York Genome Center (NYGC) (Byrska-Bishop et al. 2022). We used the phase 3 1kGP genomes only for comparison to previous releases. The phase3 1kGP dataset was downloaded from https://hgdownload.soe.ucsc.edu/gbdb/hg38/1000Genomes/. The high-coverage 1kGP genomes sequenced at the NYGC were downloaded from http://ftp.1000genomes.ebi.ac.uk/vol1/ftp/data_collections/1000G_2504_high_coverage/working/20201028_3202_raw_GT_with_annot/, which were harmonized with HGDP genomes to generate the HGDP+1kGP callset.

#### Human Genome Structural Variation Consortium (HGSVC)

The HGSVC generated high-coverage long-read WGS data and genomic variant calls from 34 samples in the 1kGP project (Ebert et al. 2021). We have evaluated precision of the SV callset by comparing against the long-read SV calls using these 34 genomes. The long-read SV calls were collected from http://ftp.1000genomes.ebi.ac.uk/vol1/ftp/data_collections/HGSVC2/release/v1.0/integrated_callset/.

#### Genome Aggregation Database (gnomAD)

We compared the HGDP+1kGP resource to gnomAD v3.1.2, which includes both HGDP and 1kGP high-coverage whole genomes, to quantify the extent of novel variation across the allele frequency spectrum contributed by these genomes. To generate allele counts and numbers in gnomAD that would be consistent with a fully non-overlapping set of genomes, we subtracted allele counts and allele numbers in the gnomAD variant callset that were contributed specifically by the 1kGP and HGDP genomes, effectively creating a synthetic version of gnomAD without these genomes.

#### Gambian Genome Variation Project (GGVP)

As part of tutorials that demonstrate how we can intersect an external dataset with HGDP+1kGP and assign meta-data labels, we intersected the HGDP+1kGP genomes with 394 Gambian Genome Variation Project genomes which are publicly available through the IGSR (http://ftp.1000genomes.ebi.ac.uk/vol1/ftp/data_collections/gambian_genome_variation_project/ <u>data</u>), as described previously (Network and Malaria Genomic Epidemiology Network 2019). We first downloaded GGVP CRAM files and used GATK HaplotypeCaller to run variant calling in GVCF mode on the 394 Gambian genome BAM files. After generating per-sample gVCFs, the single-sample gVCFs were combined into a multi-sample Hail Sparse MatrixTable (MT) using

Hail’s run_combiner() function (https://hail.is). The GGV Sparse MT was then combined with the HGDP+1kGP Sparse MT using Hail’s vcf_combiner (https://hail.is) to create a unique Sparse MT. Note that the Hail Sparse MatrixTable has since been replaced by the Hail VariantDataset.

#### Initial variant calling

The gnomAD Consortium aggregated and called variants across 153,030 individuals that included the HGDP+1kGP genomes as part of a larger project described previously (Chen et al. 2022; Karczewski et al. 2020). Briefly, using bwa mem 0.7.15.r1140, genomes were mapped to the GRCh38 version hs38DH, which includes decoy contigs and HLA genes (FASTA located at https://console.cloud.google.com/storage/browser/gcp-public-data--broad-references/hg38/v0/). Reads were then processed using GATK best practices (DePristo et al. 2011) to produce gVCFs, specifically GATK4 for all modules except HaplotypeCaller (GATK3.5). Variants from all samples in gnomAD were called jointly using the Hail combiner (Hail v0.2.62, https://hail.is) and converted to a VariantDataset (VDS), which was then densified into a dense MatrixTable used for analysis. The gnomAD team then developed and utilized an updated pipeline of sample, variant, and genotype quality control as described previously (Chen et al. 2022). We made minor modifications to these QC procedures for the extracted subset of HGDP+1kGP, as described in the sample and variant QC section.

#### Sample and variant QC

Quality control of samples was conducted according to procedures used in gnomAD, which include hard filtering with BAM-level metrics, sex inference, and ancestry inference described in greater depth previously (Chen et al. 2022). However, we modified some filtering procedures to relax some gnomAD sample QC filters new to v3 in especially diverse or unique genomes.

Specifically, the filters starting with ‘fail_’. These filters indicate whether samples are outliers in number of variants after regressing out principal components, which can indicate a sample issue. However, we identified whole continental groups and populations that were removed due solely to SNV and indel residual filters, especially those that were most genetically unique ( San, Mbuti, Biaka, Bougainville, Papuan Sepik, and Papuan Highlands). Additional individuals from the LWK, Bantu Kenya, and Bantu South Africa populations were also removed solely on the basis of the fail_n_snp_residual filter, so we removed the gnomAD ‘fail_’ filters that quantify variant count residuals after regressing out PCs.

The raw dataset includes 189,381,961 variants (SNVs and indels) and 4,150 samples. We further filtered samples and variants according to gnomAD filters. Specifically, we excluded samples that failed gnomAD’s sample QC hard filters and kept variants which were flagged as passing in the gnomAD QC pipeline. gnomAD applied an allele-specific version of GATK Variant Quality Score Recalibration (VQSR) trained on the following allele-specific features: FS, SOR, ReadPosRankSum, MQRankSum, and QC for SNPs and indels, as well as MQ for SNPs. In addition to filtering on VQSR PASS status and gnomAD sample QC filters, we also applied genotype QC filters using a function imported from gnomAD, as described previously (Chen et al. 2022), removed two contaminated samples, and removed monomorphic variants. This reduced the number of variants to 159,339,147 and removed 33 samples in total. As part of our QC process, we also updated HGDP population labels and some geographical coordinates as recommended previously (Bergström et al. 2020; Byrska-Bishop et al. 2022).

Following QC, we conducted global and subcontinental Principal Component Analysis (PCA) within and among meta-data geographical/genetic region labels (AFR, AMR, CSA, EAS, EUR, MID, and OCE) and identified 23 ancestry outliers who deviated substantially in PC space from others with the same meta-data label along the first 10 PCs (these were identified visually when one to a few individuals defined the entire PC). After removing those individuals, 153,894,851 SNVs and indels in 4,094 individuals remained.

We calculated per-sample QC metrics such as the number of SNVs and call rate using the sample_qc() method in Hail. Because singletons are especially sensitive to variation in sample size per population which is substantial across HGDP and 1kGP, we compared singleton counts by randomly downsampling to 4 unrelated samples, the minimum number of unrelated individuals per population, then removed monomorphic variants. We computed coverage data using the bam metrics field from gnomAD. We then calculated the mean of these metrics per individual within a population using Hail’s hl.agg.stats() method (https://hail.is).

#### Relatedness

We computed relatedness among 4,117 samples using the KING-robust algorithm (Manichaikul et al. 2010) implemented in Hail (https://hail.is). Specifically, we considered SNVs with MAF between 0.05 and 0.95 and missingness < 0.1%, performed LD pruning within a 500kb window, restricting to variants with r^2^ < 0.1. Then, for sample-pairs with kinship greater than 0.05, we restricted to a maximally independent set of unrelated individuals. This resulted in 200,403 SNVs and partitioned the dataset into 3,419 unrelateds and 698 relateds for PCA analysis. .

#### PCA and ADMIXTURE

We computed 20 PCs across global populations as well as within each continental ancestry group according to the “Genetic.region” project meta-data label harmonized across HGDP and 1kGP as shown in **Table S1**. After computing relatedness on the QC’ed, filtered, and LD pruned dataset as described above, we ran PCA both globally and within meta-data labels (AFR, AMR, CSA, EAS, EUR, MID, and OCE) in unrelated individuals using Hail’s hwe_normalized_pca() function (https://hail.is). We then projected related individuals into that PC space using a pc_project() function used in gnomAD and implemented in Hail. PC plots were then generated using R.

The filtered dataset was also used to run ADMIXTURE (Alexander et al. 2009) across populations and geographical regions for values of K=2 through K=10 using the command ‘admixture {bed_file} {1-10}’. We conducted 10 runs for each value of K and performed a 5-fold cross-validation error for the first run of each K by adding ‘--cv=5’ to the command. Pong (Behr et al. 2016) was used to visualize ADMIXTURE results. We selected K=6 as the best fit value of K based on a reduction in the rate of change of our 5-fold cross-validation as seen in **Figure S8**. The best fit value of K exhibits a low cross-validation error compared to other K values.

#### F_ST_ versus geographical distance

For each population pair that had an F_ST_ value, we calculated geographical distance using the haversine method (geosphere package in R) with the Earth’s radius of 6,371 km. This method of calculation did not account for human migration patterns so we additionally recalculated the pairwise geographical distances by incorporating five waypoints: Istanbul, Cairo, Phnom Penh, Anadyr and Prince Rupert, and set predetermined paths that go through certain waypoints depending on the geographical/genetic region to which the population pairs belong (Ramachandran et al. 2005). For example, to calculate the geographical distance between AMR and AFR populations, the path would go through Prince Rupert, Anadyr, and Cairo. In this example, the total distance between the pair would be the sum of the distances between the starting population and the first waypoint, pairs of waypoints in order (i.e. first to second, then second to third), and the third waypoint and the destination population. The distance between points was calculated using the haversine. We compared correlations between genetic divergence and geographical distance with and without waypoints using Pearson’s correlation and Mantel tests.

### Structural variants

#### Initial SV discovery and pruning

We applied GATK-SV (Collins et al. 2020), a public repository at https://github.com/broadinstitute/gatk-sv, to integrate and genotype SVs from the HGDP and 1kGP samples. Briefly, the HGDP samples were split into four equivalent sized batches, each consisting ∼190 samples, based on their initial cohort, PCR status, sex chromosome ploidy, and sequencing depth of the libraries (**Figure S3**). Raw Initial SVs were detected per sample by Manta (Chen et al. 2016), Wham (Kronenberg et al. 2015), MELT (Gardner et al. 2017), cnMOPs (Klambauer et al. 2012), and GATK-gCNV (Babadi et al. 2023) (see the “GatherSampleEvidence” and “GatherBatchEvidence” functions of GATK-SV) and then were clustered across each batch (see “ClusterBatch” function of GATK-SV) and filtered through an initial random forest machine learning model to remove potential false positive SVs (see “GenerateBatchMetrics” and “FilterBatch” functions of GATK-SV). The same methods were applied for SV discovery from the 1KGP samples, with details described previously (Byrska-Bishop et al. 2022). We then concatenated SVs from both HGDP and 1KGP samples to form a non-redundant set of unique SV sites (see “MergeBatchSites” function of GATK-SV), and genotyped them across all HGDP + 1KGP samples (see “GenotypeBatch” function of GATK-SV). Overlapping SVs that indicate potential complex formats of SVs were clustered and resolved into complex events (see “MakeCohortVcf” function of GATK-SV). We observed mosaicism resulting from gain or loss of X and Y chromosomes for several samples (**Table S5**), likely due to a cell line artifact from passaging. While mosaic loss of the Y chromosome is the most common form of clonal mosaicism (Thompson et al. 2019), the non-canonical sex chromosome ploidies observed are not unique to these samples and have been previously observed in other datasets (Collins et al. 2020; Byrska-Bishop et al. 2022).

#### SV refinement and annotation

A series of refinements have been applied to improve the precision of SV calls while maintaining high sensitivity. First, two machine learning models have been developed and applied to prune false positive SVs. A lightGBM model (Byrska-Bishop et al. 2022) has been trained on the 9 1kGP samples that have been deep sequenced with long-read WGS data by the HGSVC (Chaisson et al. 2019; Ebert et al. 2021), and applied to all SVs except for large bi-allelic CNVs (>5Kb). Meanwhile, a minGQ model (Collins et al. 2020) has been trained using the inheritance information among trio families to filter bi-allelic CNVs that are 5Kb and above. Genomes that failed the machine learning models were assigned a null genotype, and the proportion of null genotypes among all samples were calculated as an “no call rate” (NCR) score. SV sites that have a 10% or higher NCR were labeled as low quality variants and removed from further analyses. Then, we examined the distribution of SVs per genome to identify potential outlier samples that carry significantly more SVs than average, and also compared the frequency of SVs across each batch to identify SVs that showed significant bias (i.e. batch effects). The resulting SV callset were annotated with their frequency by their ancestry.

### Dataset Comparisons

All of the comparison datasets used GRCh38 as their reference genome except for phase 3 1kGP, which was on hg19. Phase 3 1kGP phased haplotypes on GRCh38 were provided by the International Genome Sample resource (IGSR).The comparison datasets consisted of phase 3 of the 1kGP (1000 Genomes Project Consortium et al. 2015), gnomAD v3.1.2 (Chen et al. 2022), high coverage HGDP whole genome sequences (Bergström et al. 2020), and the New York Genome Center (NYGC) 1kGP (Byrska-Bishop et al. 2022). All of these datasets were sequenced to high coverage (30X+) aside from the phase 3 1kGP, which is an integrated callset comprised of array, exome, and whole genome data with the whole genomes sequenced to 4-8X coverage. The NYGC 1kGP dataset includes all of the original 2,504 samples from phase3 1kGP as well as an additional 698 related samples. The NYGC dataset and the Bergström HGDP dataset were the only two datasets which contained multi-allelic variants, and multi-allelic variants were split before conducting the comparison. The synthetic gnomAD dataset was generated by filtering to samples only included in the gnomAD release, excluding samples which were in HGDP+1kGP but not in gnomAD. The allele count, allele number, and MAF were then calculated on this synthetic gnomAD dataset to obtain those metrics for samples which were only in gnomAD. The variant annotations for the X and Y chromosomes for the NYGC 1kGP and the Bergstrom HGDP datasets differed from those of the autosomes. Because this prevented them from being read into Hail, we removed the X and Y chromosomes from all datasets prior to conducting the comparison. In addition, each of the comparison datasets were filtered using the INFO section of their respective VCFs as imputed by the hail import_vcf() method (https://hail.is). For Bergström HGDP, those filters were excess heterozygosity (ExcHet) and LOW_VQSLOD which removed 4,278,530 total variants. NYGC 1kGP has VQSRTrancheINDEL99.00to100.00 and VQSRTrancheSNP99.80to100.00 filters which remove 10,909,291 variants. Phase 3 1kGP did not have any filters listed in the info field. The gnomAD dataset had 115,034,289 total variants removed by the following filtering: allele count is zero after filtering out low-confidence genotypes (AC0), failed Allele-Specific Variant Quality Score Recalibration (AS_VQSR), and inbreeding coefficient< -0.3 (InbreedingCoeff). Counts for the number of variants removed by comparison dataset filters can be found in **Supplementary Table S10**. Variant comparison was done using MAF which was calculated for each comparison dataset apart from gnomAD using Hail’s variant_qc() method (https://hail.is) and taking the minimum value of the allele frequency array. A MAF table was created for each of the comparison datasets containing counts of the number of variants in each MAF bin. To generate these tables, we first removed any missing variants. We then created a flag, in_comparison, which was True if a variant in the comparison dataset at [locus, alleles], was also in the HGDP+1kGP dataset. The HGDP+1kGP dataset used for every comparison aside from gnomAD was the post-QC version. This version excludes PCA outliers and has gnomAD sample, variant, and genotype QC applied, as described in the Sample and variant QC section. Due to the synthetic version of the gnomAD dataset used for comparison, the post-QC HGDP+1kGP comparison dataset included PCA outliers, as some of them may have been in gnomAD. MAF bins were created containing the counts of variants from HGDP+1kGP in each of the MAF categories. Once the bins were generated, we used the in_comparison flag to see which alleles in the dataset were shared and in HGDP+1kGP only. In the comparison dataset we created a flag, not_in_hgdp_1kg, which was True if the variants in the comparison dataset were not present in HGDP+1kGP. We appended the count of True values for that flag onto the table as the 0% MAF bin, to denote variants which were only in the comparison dataset and therefore have a MAF of 0% in HGDP+1kGP. The final bar plot was generated from the data in the MAF tables using R. To investigate why some variants were missing in the HGDP+1kGP dataset compared to the comprising datasets, we looked at pre- and post-QC counts of SNPs and indels for these variants, finding that there are less variants missing from HGDP+1kGP after quality control filters were applied to the comparison dataset (**Supplementary Table S11**). For all of the datasets aside from NYGC 1kGP, we used the included annotations of SNV or indel to calculate the numbers. NYGC 1kGP did not have allele type information, so we manually classified indel and SNVs based on length of alleles. The phase3 1kGP phased dataset did not contain any indels. For the gnomAD dataset there were additionally variants labeled as complex, which we excluded from these counts. Alleles labeled as deletion and insertion were combined to create one indel label. To calculate the SNV and indel counts, we took the intersection of variants which were in the comparison dataset but not in the HGDP+1kGP dataset, as with the main comparison described above, additionally counting variants in pre-and post-QC datasets. We did this 4 times to obtain all combinations of pairwise pre- and post-QC comparisons, then counted the number of variants in the pre- and post-QC HGDP+1kGP dataset.

### Phased haplotypes and imputation accuracy evaluation

To create a haplotype reference panel that can be used for phasing or imputation, we first constructed a pedigree file with familial relationships between first degree relatives in the quality controlled harmonized dataset to improve phasing accuracy. We ran additional relatedness checks using the PC-Relate algorithm (Conomos et al. 2016) implemented in Hail (https://hail.is). The PC-Relate results were filtered to sample pairs with a kinship statistic between 0.248 and 0.252. We then cross-checked the filtered PC-Relate results with the publicly available NYGC 1kGP pedigree file (Byrska-Bishop et al. 2022) and found that all parent-child relationships estimated by PC-Relate are reported in the 1kGP pedigree file. Of 602 previously reported trios, 9 samples failed QC (6 due to gnomAD sample QC, 3 due to ancestry outliers). In total, we therefore included 599 families, 6 of which were duos and the remaining 593 of which were trios. To investigate if there are any possible duplicate samples/monozygotic twins within or across projects, we filtered the PC-Relate results to sample pairs with kin statistic > 0.35 and found 5 pairs of samples, of which 3 have been reported before (Mountain and Ramakrishnan 2005). To verify if the 5 sample-pairs are indeed possible duplicates and/or monozygotic twins, we ran Identity-By-Descent (ref) as implemented in Hail (https://hail.is) and found that each sample-pair shared almost all alleles (IBS2). One sample from each of the 5 pairs was filtered out from the dataset.

As recommended by the SHAPEIT5 documentation, we applied additional QC filters to the dataset before phasing the haplotypes, keeping only variants with: (1) HWE>=1E-30; (2) F_MISSING<=0.1; and (3) ExcHet>=0.5 && ExcHet<=1.5. Common variants (MAF >= 0.1%) were phased in large chunks of length 20cM using the phase_common program in SHAPEIT5. The common variants chunks were then ligated together to create a haplotype scaffold containing partially phased haplotypes for each autosome (chr1-22). Using the partially phased scaffolds as input, rare variants (MAF < 0.1%) were then phased in small chunks of length 4cM using the phase_rare program in SHAPEIT5. Lastly, the fully phased chunks were then concatenated into chromosomes and indexed using bcftools. To improve the quality of phasing, pedigree information was used when phasing both common and rare variants.

To evaluate phasing accuracy, we used the “switch” utility of SHAPEIT5 to compute switch error rate, treating the HGSVC2 genomes as “truth” and statistically phased genomes as a test set. To evaluate imputation accuracy, we used a filtering and downsampling strategy to simulate GWAS arrays and 93 whole genomes sequenced at various depths from previously sequenced high-coverage whole genomes from the Neuropsychiatric Genetics of African Populations-Psychosis (NeuroGAP-Psychosis) study, as previously (Martin et al. 2021). Briefly, these genomes were from participants in Ethiopia, Kenya, South Africa, and Uganda. Ethical and safety considerations are being taken across multiple levels, as described in greater detail previously (Stevenson et al. 2019). The GWAS arrays we evaluated included the widely used Illumina Global Screening Array (GSA) designed to increase scalability and improve imputation accuracy in European populations, the Multi-ethnic Genotyping Array (MEGA) designed to improve performance across globally diverse populations, and H3Africa array specialized for higher genetic diversity and smaller haplotype blocks in African genomes. We previously downsampled reads randomly to an average of 0.5X, 1X, 2X, and 4X using the GATK DownsampleSam module, which retains a random subset of reads and their mate pairs deterministically. More details on the downsampling strategy are in (Martin et al. 2021).

## Code

https://github.com/atgu/hgdp_tgp

## Data availability

All data are freely available and described in more detail here: https://gnomad.broadinstitute.org/news/2020-10-gnomad-v3-1-new-content-methods-annotations-and-data-availability/#the-gnomad-hgdp-and-1000-genomes-callset.

The gnomAD HGDP+1kGP callset can be found here: https://gnomad.broadinstitute.org/downloads#v3-hgdp-1kg

Computational tutorials for conducting many of the most common genetic analyses including those implemented on this dataset are available here: https://github.com/atgu/hgdp_tgp/tree/master/tutorials

Datasets used in the tutorials are located here: https://gnomad.broadinstitute.org/downloads#v3-hgdp-1kg-tutorials

Phased haplotypes are available as BCFs on google cloud at the following path: gs://gcp-public-data--gnomad/resources/hgdp_1kg/phased_haplotypes_v2/ More details can be found here: https://docs.google.com/document/d/1LCx74zREJaJwtN0MzonSv1QB3UahVtgTfjkepXaQUxc/edit.

These datasets are released on Google Cloud Platform, Amazon Web Services, and Microsoft Azure, and can be found on the Downloads page of the gnomAD browser (https://gnomad.broadinstitute.org/downloads#v3-hgdp-1kg). Instructions on how to download these datasets can be found here: https://gnomad.broadinstitute.org/downloads.

## Author contributions

Z.K., M.T.Y., L.L.N., H.A.K., A.R.M. performed analysis. J.K.G., M.W.W., K.R.C., G.T., and K.J.K. processed these data with gnomAD. X.Z., S.P.H., M.A.W., M.E.T., and H.B. called structural variants. M.J.D., E.G.A., and A.R.M. conceptualized the project. A.R.M. provided funding. Z.K., M.T.K., K.J.K., E.G.A., and A.R.M. wrote the paper with input from all co-authors.

## Acknowledgements

This work was supported by funding from the National Institutes of Health (K99/R00MH117229 to A.R.M., R01DE031261 to H.B. and A.R.M., R01MH115957 to M.E.T.). This project was also supported by funding from BroadIgnite, Broad Next Generation Fund, and a Merkin Fellowship Award.

