## Supplemental Figures 1-20, Tables 1-13 for "A harmonized public resource of deeply sequenced diverse human genomes"

### Supplemental materials

**Supplemental materials** **1**
Data harmonization 3
Genotype data 3
Meta-data 3
Table S1 | Harmonization of HGDP and 1000 Genomes Project meta-data project labels. 3
Table S2 | Genetic outliers identified in analysis of global and subcontinental PCA. 4
Table S3 | Final sample counts. 5
QC Meta-data Summaries 5
Figure S1 | Coverage across the 1kGP and HGDP. 5
Figure S2 | Coverage across 1kGP and HGDP by population. 6
Table S4 | Coverage as well as SNV and SV statistics by population. 6
Structural variants (SVs) 7
Figure S3 | Dosage and sex ploidy of HGDP samples and batching strategy. 7
Figure S4 | SV callset and quality evaluation results. 8
Table S5 | Sex chromosome aneuploidies in the HGDP samples. 8
Figure S5 | Mean count of SVs versus SNVs by project, region, and number of individuals. 9
Table S6 | SV calls by external support from the HGSV study. 10
Figure S6 | SV breakdown in count by class across HGDP and 1kGP (HGSV). 11
Population genetic comparisons 12
Figure S7 | ADMIXTURE analysis of the HGDP and 1kGP resource. 13
Figure S8 | 5-fold cross-validation error across ADMIXTURE runs. 14
Figure S9 | PCA biplots and densities globally. 15
Figure S10 | Subcontinental PCA in AFR populations. 15
Figure S11 | Subcontinental PCA in CSA populations. 16
Figure S12 | Subcontinental PCA in EAS populations. 16
Figure S13 | Subcontinental PCA in EUR populations. 17
Figure S14 | Subcontinental PCA in AMR populations. 17
Figure S15 | Subcontinental PCA in MID populations. 18
Figure S16 | Subcontinental PCA in OCE populations. 18
Figure S17 | HGDP+1kGP ancestry labels applied to the Gambian Genome Variation (GGV) Project. 19
Figure S18 | Dendrogram of the pairwise FST heatmap between populations colored by geographical/genetic regions. 20
Table S7 | Populations interspersed among other geographical/genetic regions obtained from the pairwise FST heatmap, colored by region. 21
Table S8 | Pearson's correlation and Mantel tests results with and without waypoints. 22
Quality control 23
Figure S19 | Example of a filter that was included in gnomAD v3.1 but excluded from this project. 24
Table S9 | gnomAD Sample and Variant Quality Control Steps 25
Dataset Comparisons 26
Table S10 | Filters Applied to Comparison Datasets 27

Table S11 | Variants absent in HGDP+1kGP that were identified in comparison datasets depending on whether variants passed QC in each dataset. 27
Table S12 | Variant counts and percentages of comparison datasets. 28
Analysis tutorials 30
Figure S20 | PCA shrinkage analysis to determine acceptable levels of missingness before ancestry resolution becomes too low to accurately assign population labels. 30
Table S13 | Shrinkage analysis matches and no classification numbers by SNP missingness in the test dataset, as shown in Figure S20. 31
References 31

### Data harmonization

#### Genotype data

Genotype data was processed as described in (Chen et al. 2022). Briefly, reads were mapped using BWA-MEM, then filtered using the GATK Best Practices pipeline, and gVCFs were generated using GATK HaplotypeCaller. Joint calling was performed using the Hail combiner (Hail Team 2021) and converted to a VariantDataset (VDS), which was then densified into a dense MatrixTable used for analysis. These datasets are released on Google Cloud Platform, Amazon Web Services, and Microsoft Azure, and can be found on the Downloads page of the gnomAD browser (https://gnomad.broadinstitute.org/downloads#v3-hgdp-1kg).

#### Meta-data

Where possible, we combined meta-data from the 1000 Genomes Project and HGDP by combining the "super population" data from the 1000 Genomes project (1000 Genomes Project Consortium et al. 2015) and region information from HGDP (Bergström et al. 2020). We created a harmonized combined label with 3-letter codes for all groups, which we refer to as geographical/genetic regions throughout the text. Where a region was only clearly contained in HGDP, we used the HGDP information to define a 3-letter code. The CENTRAL_SOUTH_ASIA code contained within HGDP is more geographically expansive than the SAS label contained in the 1000 Genomes Project, so we expanded the 3 letter code to be CSA, as shown in **Table S1**.

**Table S1 | Harmonization of HGDP and 1000 Genomes Project meta-data project labels.**
These labels are referred to as geographical/genetic regions throughout this manuscript. The sample sizes included here are pre-QC. Post-QC numbers are shown in **Table S3**.

| 1000 Genomes super population | HGDP region | Combined label (geographical/ genetic region) | Sample size (pre-QC) |
|---|---|---|---|
| AFR | AFRICA | AFR | 1003 |
| AMR | AMERICA | AMR | 552 |
| SAS | CENTRAL_SOUTH_ASIA | CSA | 790 |
| EAS | EAST_ASIA | EAS | 825 |
| EUR | EUROPE | EUR | 788 |
| N/A | MIDDLE_EAST | MID | 162 |
| N/A | OCEANIA | OCE | 30 |

After combining region data, we used principal components analysis (PCA) to identify ancestry outliers within regions. We identified outliers as described in **Table S2** and provide final sample counts in **Table S3**.

**Table S2 | Genetic outliers identified in analysis of global and subcontinental PCA.**
Within subcontinental PCA biplots, outliers were identified when one to three samples defined most of an entire PC among PCs 1-6.

| Sample ID | Region | Population |
|---|---|---|
| HG01880 | AFR | ACB |
| HG01881 | AFR | ACB |
| NA20274 | AFR | ASW |
| NA20299 | AFR | ASW |
| NA20314 | AFR | ASW |
| HGDP00013 | CSA | Brahui |
| HGDP00029 | CSA | Brahui |
| HGDP00057 | CSA | Balochi |
| HGDP00130 | CSA | Makrani |
| HGDP00150 | CSA | Makrani |
| HGDP00175 | CSA | Sindhi |
| HGDP01298 | EAS | Uygur |
| HGDP01300 | EAS | Uygur |
| HGDP01303 | EAS | Uygur |
| LP6005443-DNA_B02 | EAS | Uygur |
| HG01628 | EUR | IBS |
| HG01629 | EUR | IBS |
| HG01630 | EUR | IBS |
| HG01694 | EUR | IBS |
| HG01696 | EUR | IBS |
| HGDP00621 | MID | Bedouin |
| HGDP01270 | MID | Mozabite |
| HGDP01271 | MID | Mozabite |

**Table S3 | Final sample counts.**
Note: hard filtering was performed as in gnomAD v3 with modifications as described below in Quality Control. The first row of the “Total” column includes a “synthetic diploid” QC sample (CHM; Complete Hydatidiform Mole) described previously (Li et al. 2018). It was removed during initial QC together with 31 samples that failed gnomAD's sample QC hard filters and two contaminated samples, and is excluded from the sample count of the initial dataset reported in the manuscript.

| | HGDP | 1kGP | Total |
|---|---|---|---|
| Initial dataset | 948 | 3,202 | 4,151 |
| Hard filtered | 941 | 3,176 | 4,117 |
| PCA outliers removed | 928 | 3,166 | 4,094 |
| Unrelated individuals | 880 | 2,520 | 3,400 |

### QC Meta-data Summaries

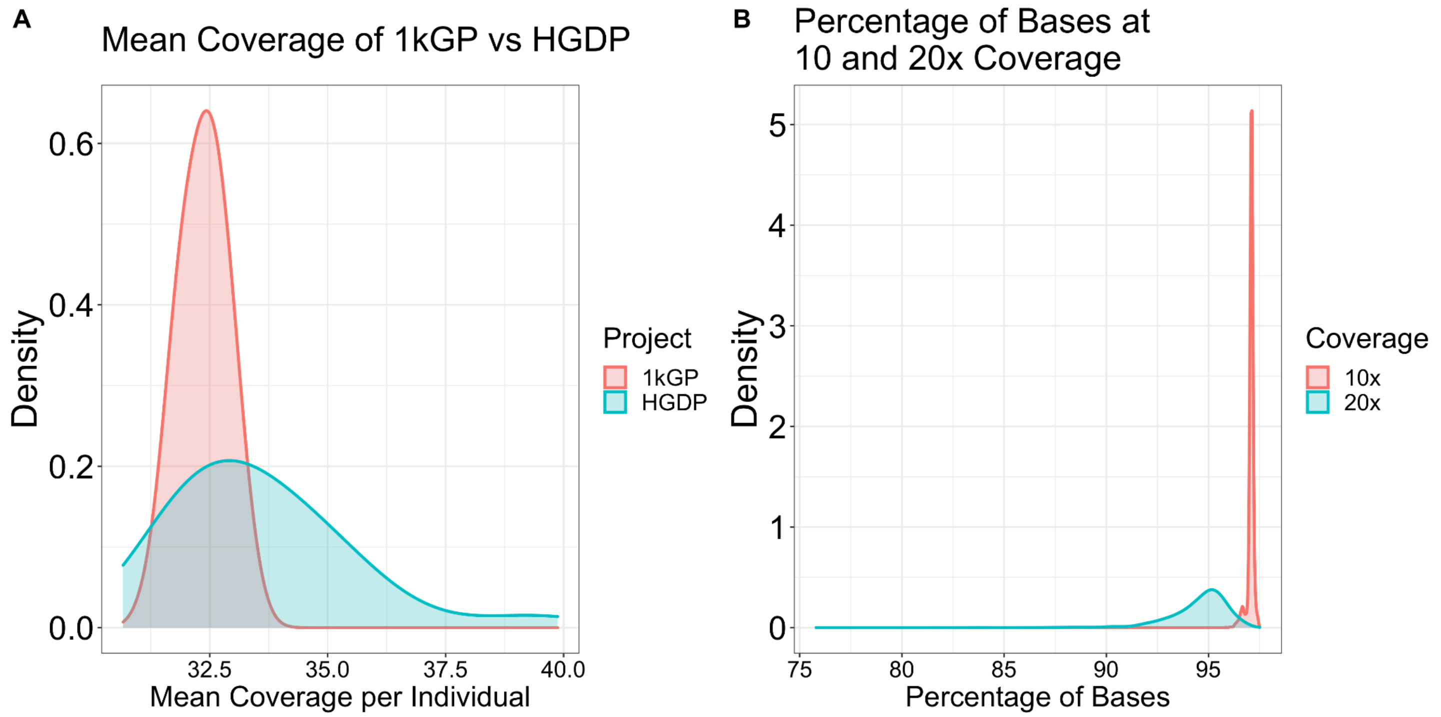


**Figure S1 | Coverage across the 1kGP and HGDP.**
A) Coverage in both datasets is uniformly above 30X, with an average of 33X coverage across the harmonized dataset. The coverage of the HGDP genomes is more variable than in 1kGP, as expected based on a variety of technical differences such as multiple sequencing batches, PCR+ vs PCR-free, and older cell lines in HGDP compared to 1kGP. The differences in project coverages also impacts the distribution of coverage statistics by Geographical region given their tally by project (**Table S4**). The overall coverage distributions by population are shown in **Figure S2**. B) Over 95% of bases are covered over 10X, and over 90% of bases are covered over 20X in HGDP+1kGP.

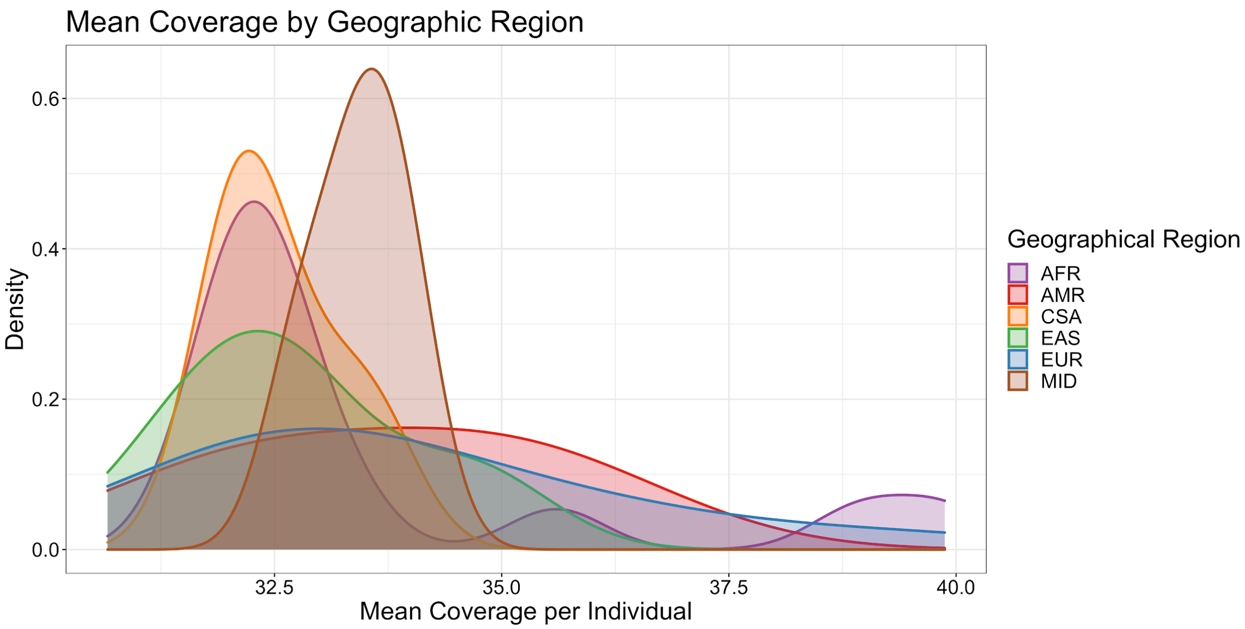


**Figure S2 | Coverage across 1kGP and HGDP by population.**
Regional abbreviations are as described in **Table S1**. OCE is excluded from this plot as it is represented by only two populations. Mean coverage across the different regions is 33X with coverage consistently above 30X for all regions.

**Table S4 | Coverage as well as SNV and SV statistics by population.**
Coverage was computed across the genome as part of the gnomAD project. Relatedness was inferred using KING-robust. Because number of variants and singleton counts per individual are sensitive to sample size imbalances, they were tallied using a downsampled version of the dataset in which each population was randomly downsampled to match the smallest population (i.e. 6 individuals per population), then SNVs were removed if they were not polymorphic in the downsampled dataset. Given the more pronounced impact of batch effects on structure variant (SV) calling and the number of batches present within and between datasets, the number of SVs per individual were calculated across the full dataset, not in the downsampled dataset.

#### Structural variants (SVs)

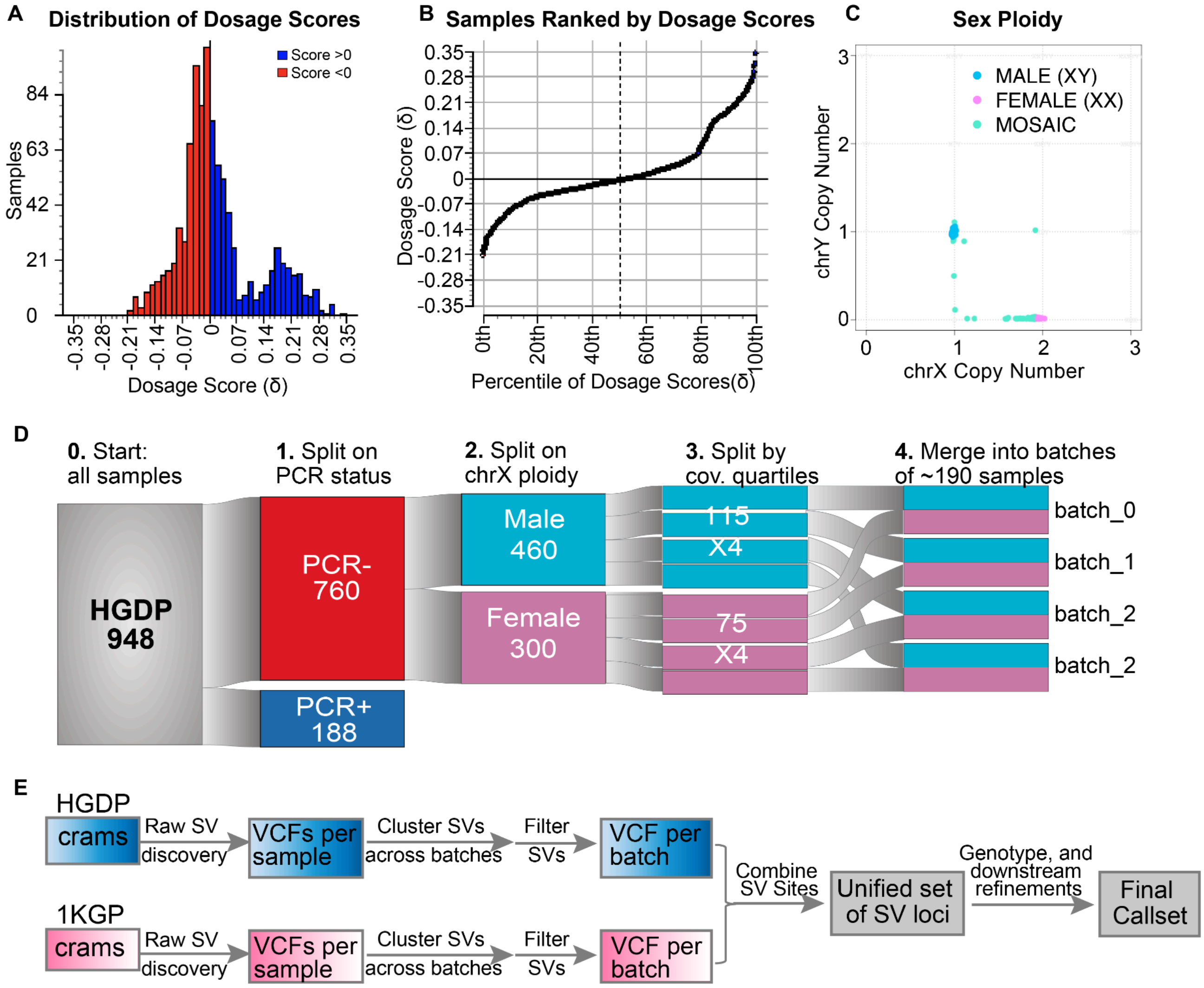


**Figure S3 | Dosage and sex ploidy of HGDP samples and batching strategy.**
A) Distribution of dosage scores across HGDP samples. We used the previously developed whole genome dosage model (Collins et al 2020) to quantify non-uniform distribution of sequencing coverage.The dosage scores corresponded predominantly to PCR-amplified (PCR+) and PCR-free (PCR-) library protocols. B) Samples ranked by dosage score. C) Distribution of chrX copy number across HGDP samples. D) Batching strategy for SV calling. HGDP samples were first split by their PCR status and chrX ploidy. PCR- samples were then ranked by their sequencing depth from low to high, and split into four sub batches of equivalent sizes. Male and female batches with matched coverage quantiles are combined to form the final batches. E) Workflow of SV discovery from the HGDP and 1KGP genomes. The HGDP and 1KGP samples have been processed separately through the first steps of GATK-SV, including raw SV discovery, batching SVs across each batch and initial filtering of SVs using the “FilterBatch” method in GATK-SV. The filtered SVs were then merged across HGDP and 1KGP to form a non-redundant set of SV loci, systematically genotype across both HGDP and 1KGP samples, and processed through downstream steps of GATK-SV (see https://github.com/broadinstitute/gatk-sv for details).

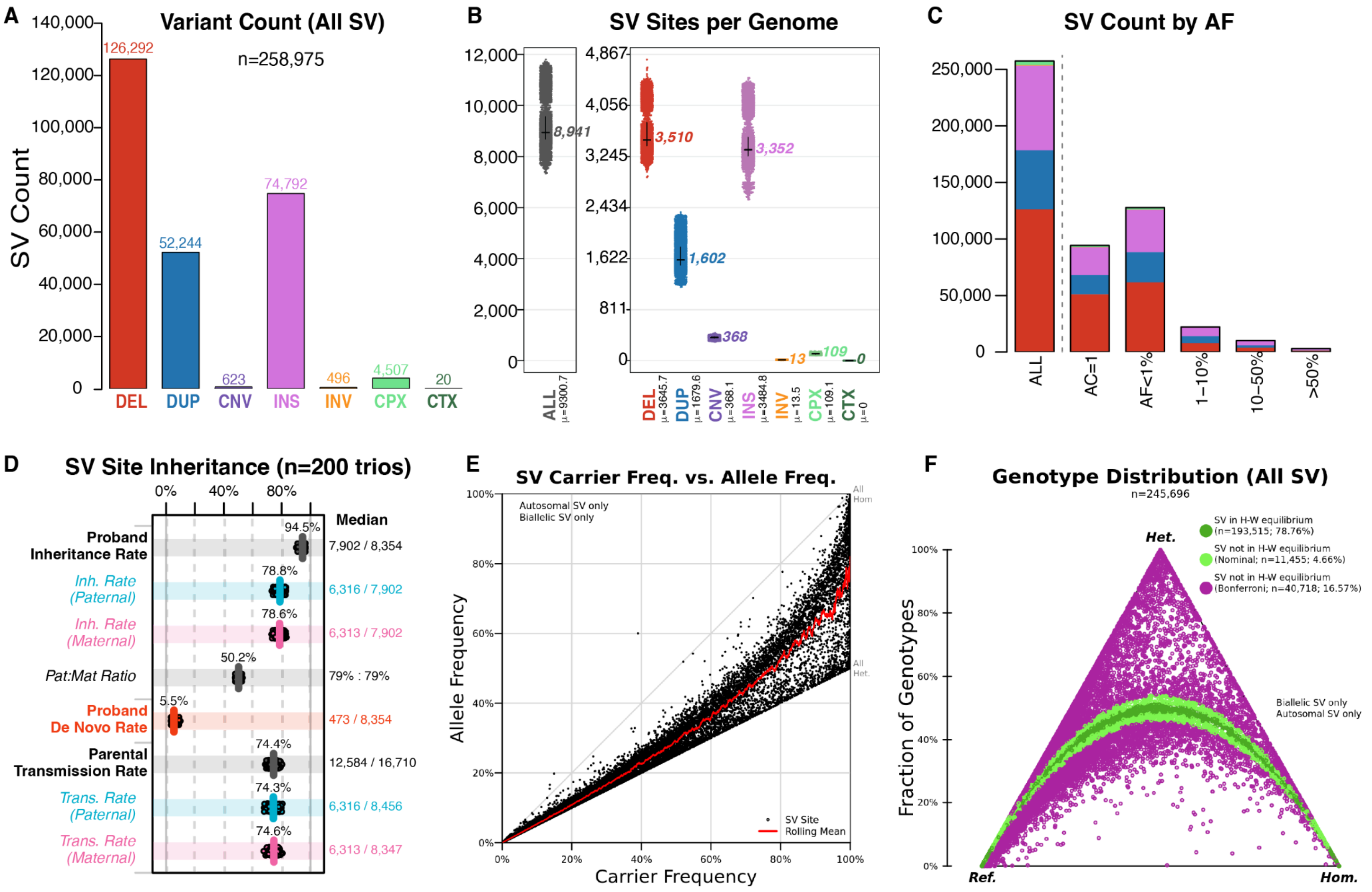


**Figure S4 | SV callset and quality evaluation results.**

A) Count of SV sites across 4,151 HGDP and 1kGP samples by variant type. B) Count of SVs per genome by variant type. C) Count of SV sites by allele frequency. D) Inheritance of SVs calculated in 100 pather-mother-child trio families. Proband Inheritance Rate - proportion of SVs in children's genome that were inherited from either parents; Paternal Inheritance Rate - proportion of SVs in children's genome that were shared by paternal genome; Maternal Inheritance Rate - proportion of SVs in children's genome that were shared by maternal genome; Parental Transmission Rate - proportion of SVs in parents' genome that were transmitted into children's genome; Trans. Rate (Paternal) - proportion of SVs in paternal genome that were transmitted into children's genome; Trans. Rate (Maternal) - proportion of SVs in maternal genome that were transmitted into children's genome. E) Correlation of allele frequencies. F) Hardy-Weinberg Equilibrium distribution of SVs across all samples. Each point is a single biallelic autosomal SV projected onto HWE ternary axes corresponding to its ratio of homozygous reference (0/0), heterozygous (0/1), and homozygous alternate (1/1) genotypes across all samples in the indicated population. The distance of a point to a vertex indicates the fraction of samples with that genotype. Deviation from HWE was assessed using a chi-square goodness-of-fit test with one degree of freedom, and points are colored based on their P-value. Green points are SVs within bounds defined for HWE based on the number of sites documented in each population, and purple points are SVs outside of these P-value bounds. The proportion of SVs corresponding to each P-value cutoff is provided at the right of each panel. Plots were generated using the "HardyWeinberg" package in R.

**Table S5 | Sex chromosome aneuploidies in the HGDP samples.**

| Sample ID | Population | Genetic region | chrX | chrY | Assignment |
|---|---|---|---|---|---|

| | | | | | |
|---|---|---|---|---|---|
| HGDP00445 | Burusho | CSA | 1 | 0 | XO |
| HGDP01157 | Bergamo Italian | EUR | 1 | 0 | XO |
| HGDP01208 | Oroquen | EAS | 2 | 1 | XXY |
| HGDP01368 | Basque | EUR | 1 | 0 | XO |
| LP6005441-DNA_G09 | Palestinian | MID | 1 | 0 | XO |

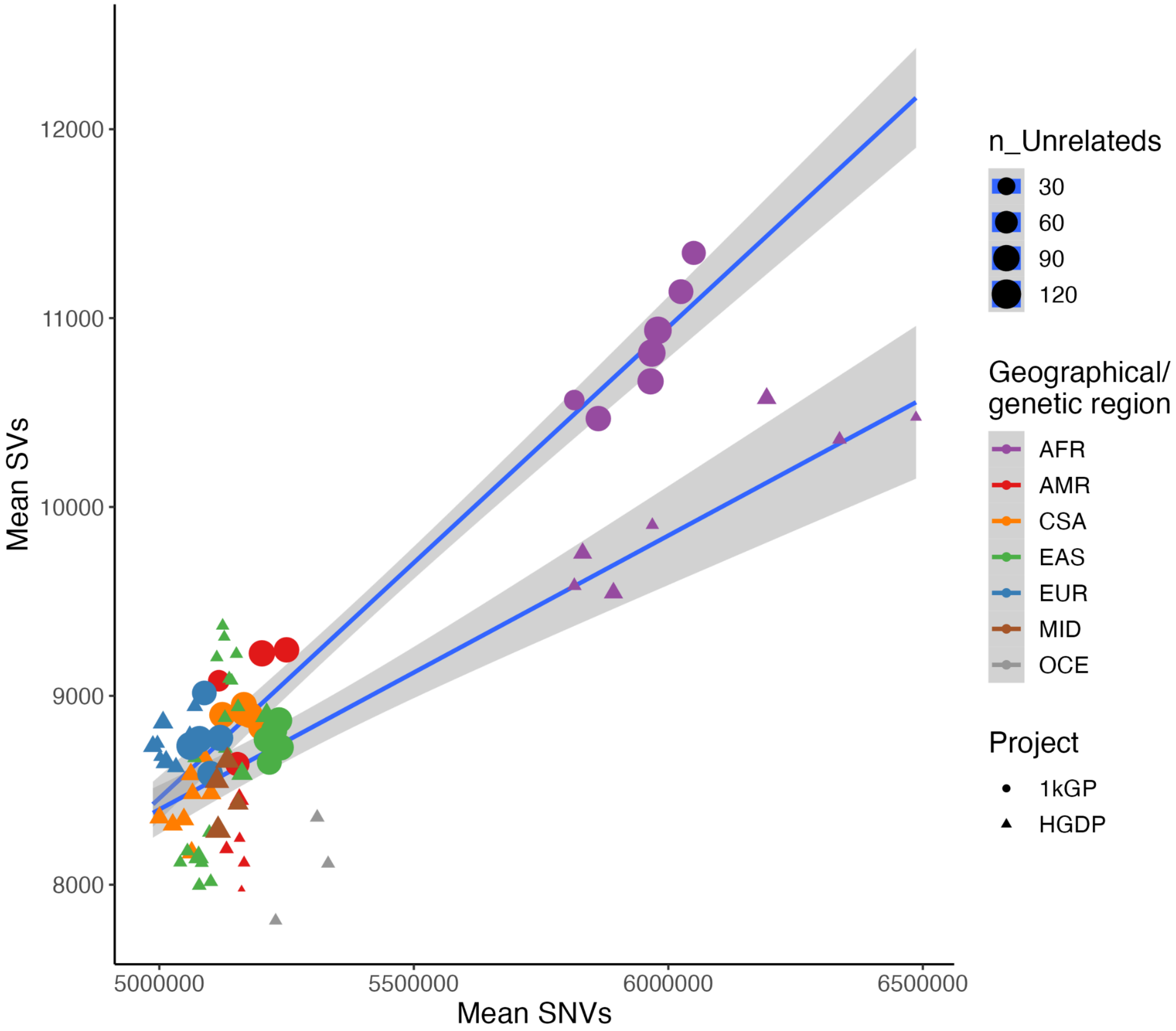


**Figure S5 | Mean count of SVs versus SNVs by project, region, and number of individuals.**
Top line shows a fitted regression line to the 1000 Genomes Project points, and bottom line is fitted to HGDP points. A larger number of SVs are present in the 1000 Genomes Project data, which was explored more fully in **Figure S6**.

**Table S6 | SV calls by external support from the HGSV study.**

| | | External Supports (Count SVs per genome) | | | |
|---|---|---|---|---|---|
| SV type | Precision | No Support | Illumina | PacBio | Illumina and PacBio |
| DEL | 96.22% | 140 | 15 | 626 | 2918 |
| DUP | 96.32% | 64 | 13 | 8512 | 808 |
| INS | 96.19% | 136 | 3 | 1049 | 2390 |
| INV | 96.62% | 1 | 1 | 0 | 13 |
| CPX | 74.54% | 28 | 3 | 134 | 64 |
| All SVs | 95.97% | 368 | 35 | 2541 | 6194 |

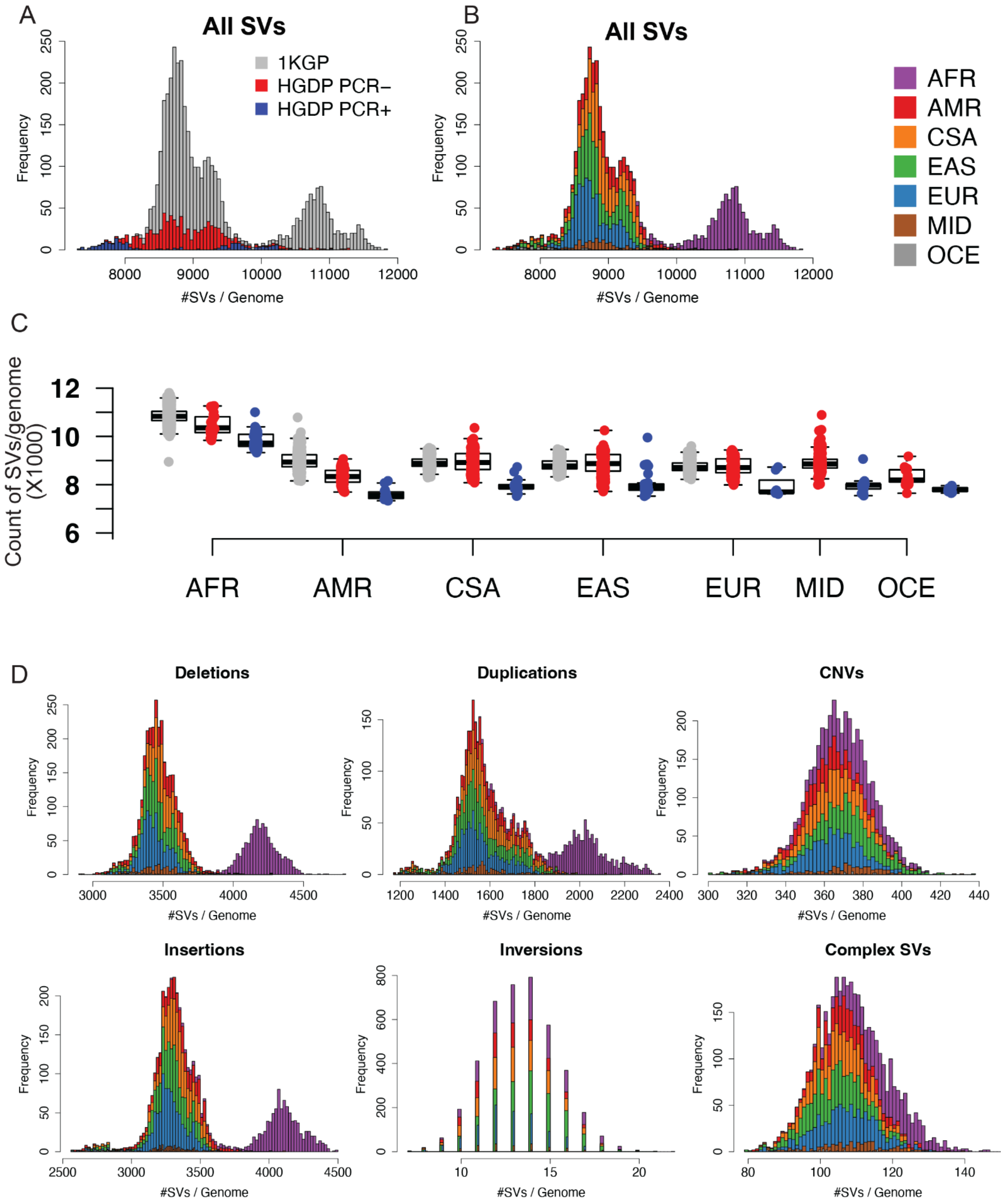


**Figure S6 | SV breakdown in count by class across HGDP and 1kGP (HGSV).**
Per genome SV counts by study and PCR status (A,C), and population (B). Per genome SV counts are also broken down by SV type, including deletions, duplications, multi-allelic CNVs, insertions, inversions, and complex SVs in D).

#### Population genetic comparisons

The breakdown of ancestry and population structure by ADMIXTURE is similar to that identified in global PCA, with K=2 highlighting structure in the AFR, K=3 highlighting structure in the EAS, K=4 highlighting structure in the EUR and CSA, K=5 highlighting structure in the AMR, K=6 highlighting structure in the OCE, K=7 highlighting structure in the MID, and subsequent values of K highlighting structure within meta-data labels (**Figure S7**).

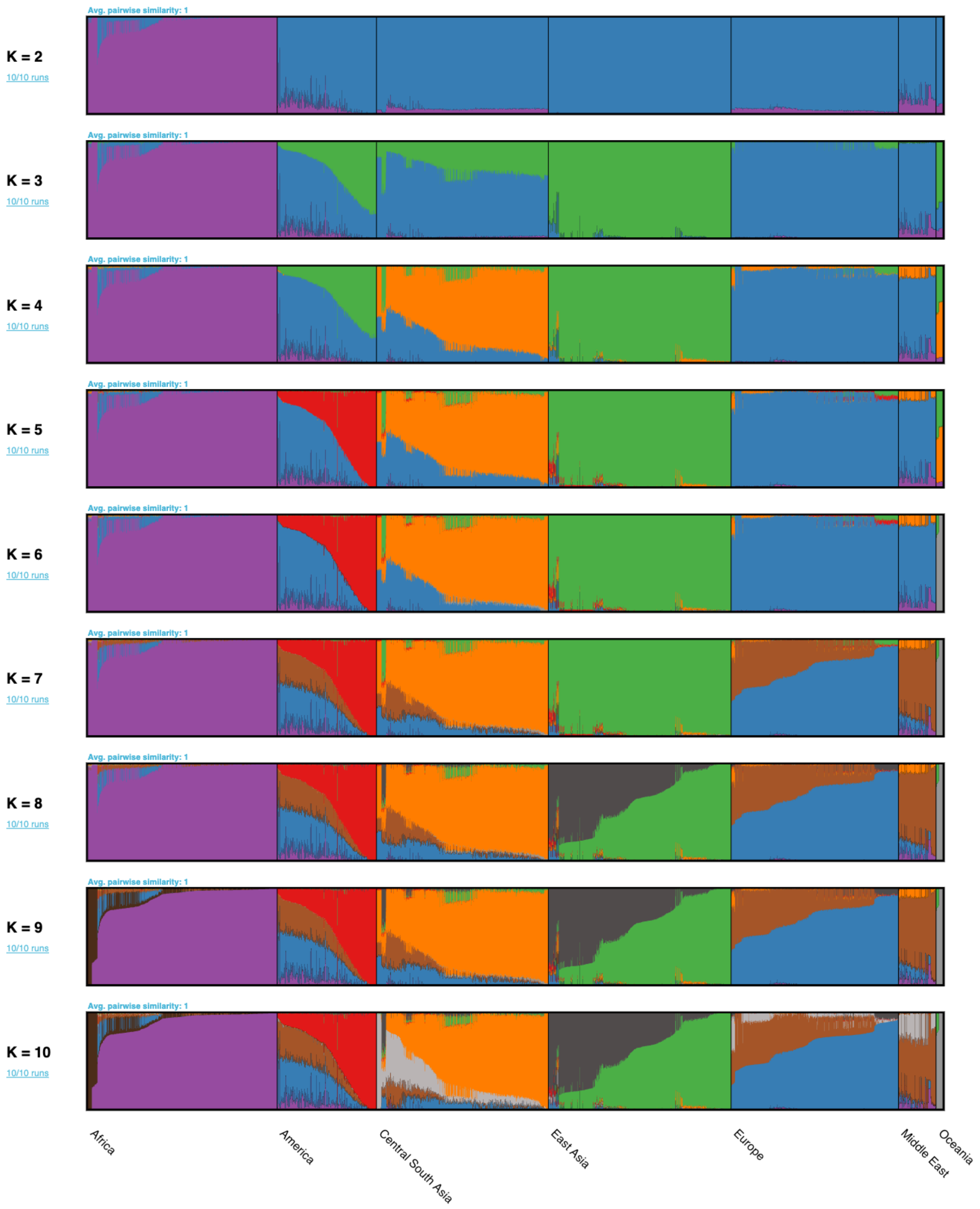


**Figure S7 | ADMIXTURE analysis of the HGDP and 1kGP resource.**
We ran ADMIXTURE with values of K=2 through K=10 across populations and harmonized geographical/genetic regions. Each row of bar plots shows the breakdown of regional substructure as K increases, where K is the number of genetic ancestry components fit in that run. For example, when K=2, AFR separates from the rest of the populations as the most distinct population due to high levels of genetic diversity.

When K=3 EUR separates from the rest, and so on. We chose the best fit value of K to be K=6 based on a reduction in the rate of change of 5-fold cross validation error as shown in **Figure S8**.

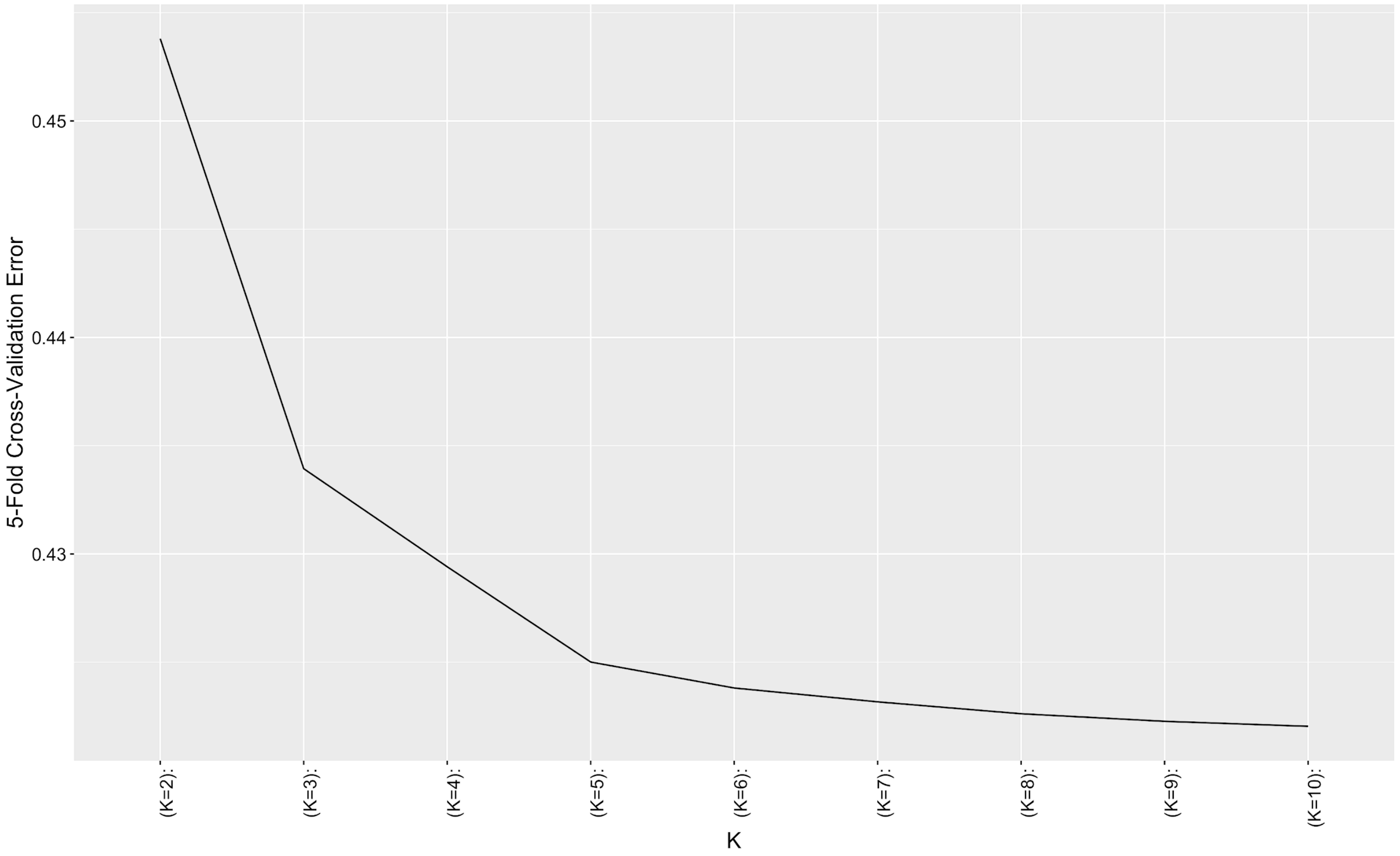


**Figure S8 | 5-fold cross-validation error across ADMIXTURE runs.**
We selected K=6 as the point at which cross-validation error leveled out. As described in the ADMIXTURE manual, the cross-validation error enables users to identify the value of K for which the model has best predictive accuracy, as determined by “holding out” data points. It partitions observed genotypes into 5 roughly equally sized folds, masks genotypes for each fold, then predicts the genotypes.

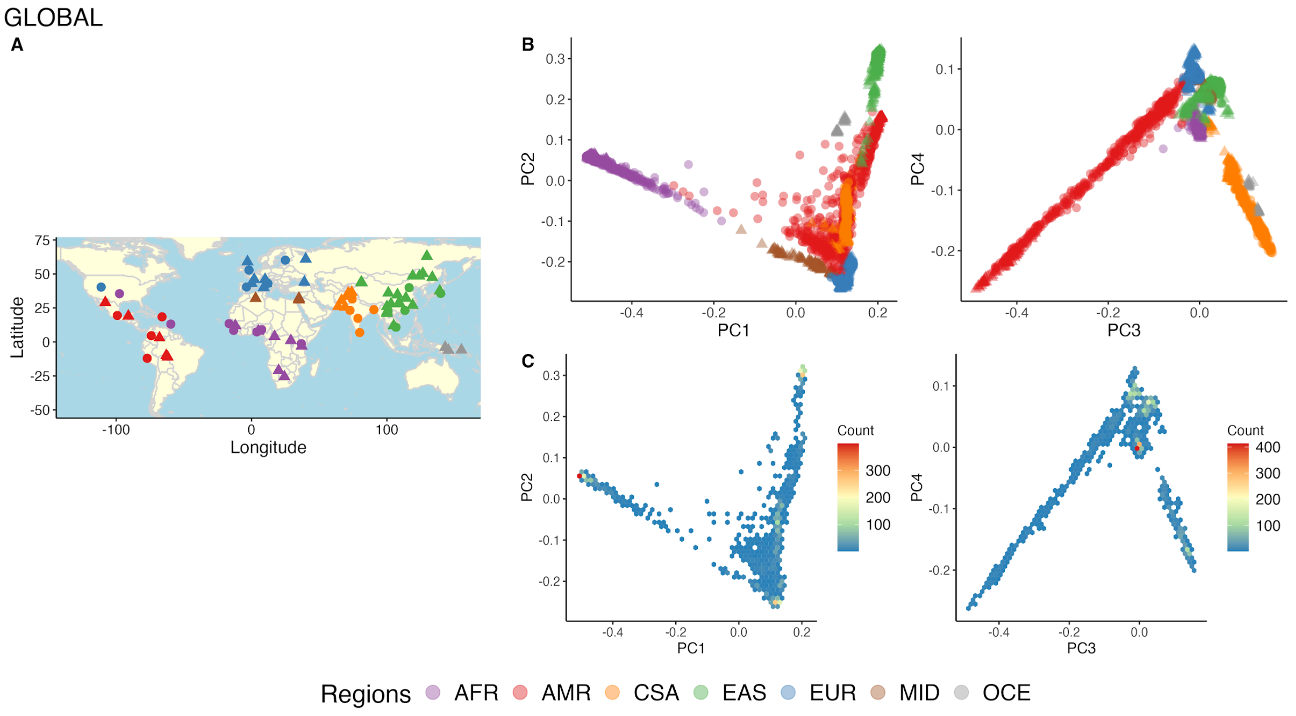


**Figure S9 | PCA biplots and densities globally.**
A) Map shows where all samples in analyses are from. B) PCA biplots of PCs 1-4. PCA outliers were removed prior to this analysis. Filled circles indicate populations in the 1000 Genomes Project, while filled triangles indicate populations in HGDP. Population codes are as in **Table S1**. C) Density plot of PCA biplots of PCs 1-4.

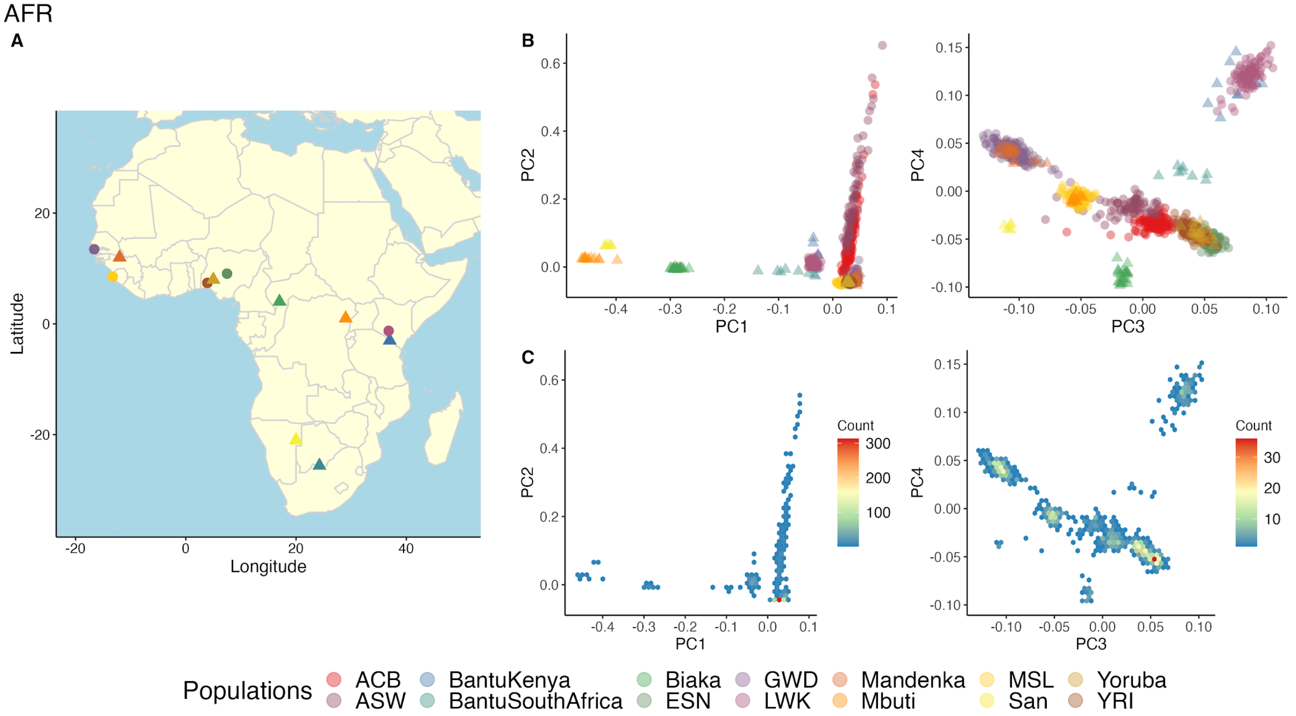


**Figure S10 | Subcontinental PCA in AFR populations.**
A) Map shows where all AFR samples in analyses are from. B) PCA biplots of PCs 1-4. PCA outliers were removed prior to this analysis. Filled circles indicate populations in the 1000 Genomes Project, while filled triangles indicate populations in HGDP. Population codes are as in **Table S1**. C) Density plot of PCA biplots of PCs 1-4.

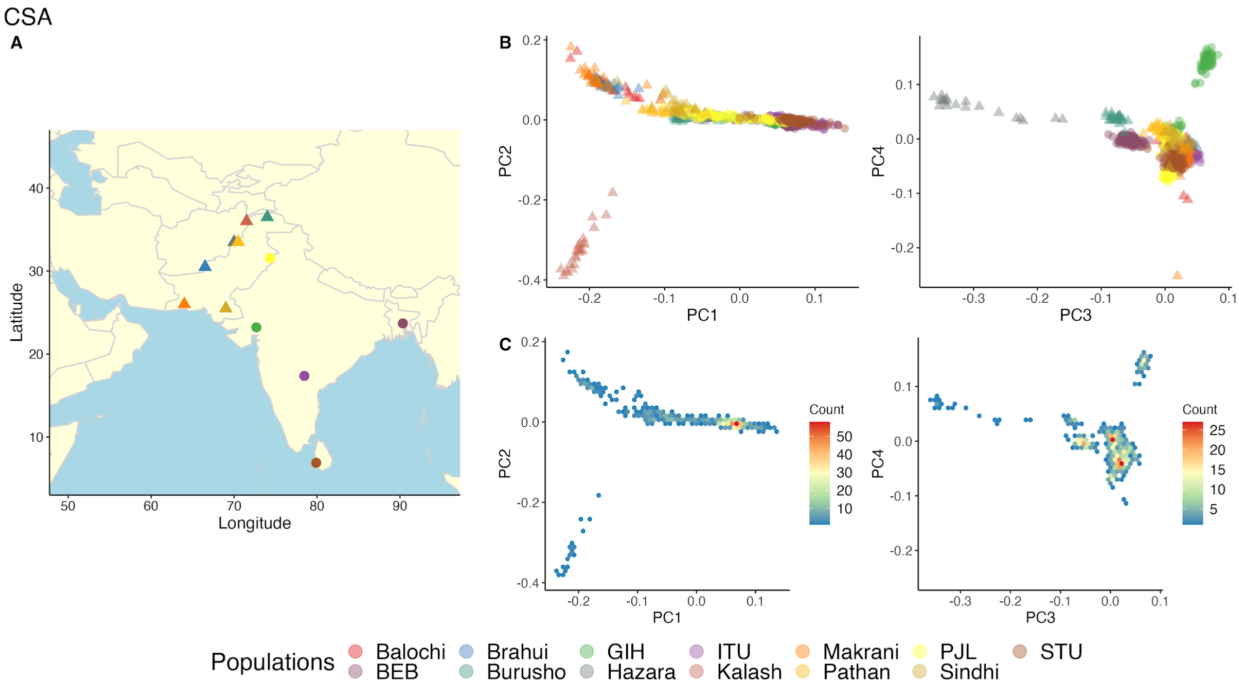


**Figure S11 | Subcontinental PCA in CSA populations.**

A) Map shows where all CSA samples in analyses are from. B) PCA biplots of PCs 1-4. PCA outliers were removed prior to this analysis. Filled circles indicate populations in the 1000 Genomes Project, while filled triangles indicate populations in HGDP. Population codes are as in **Table S1**. C) Density plot of PCA biplots of PCs 1-4.

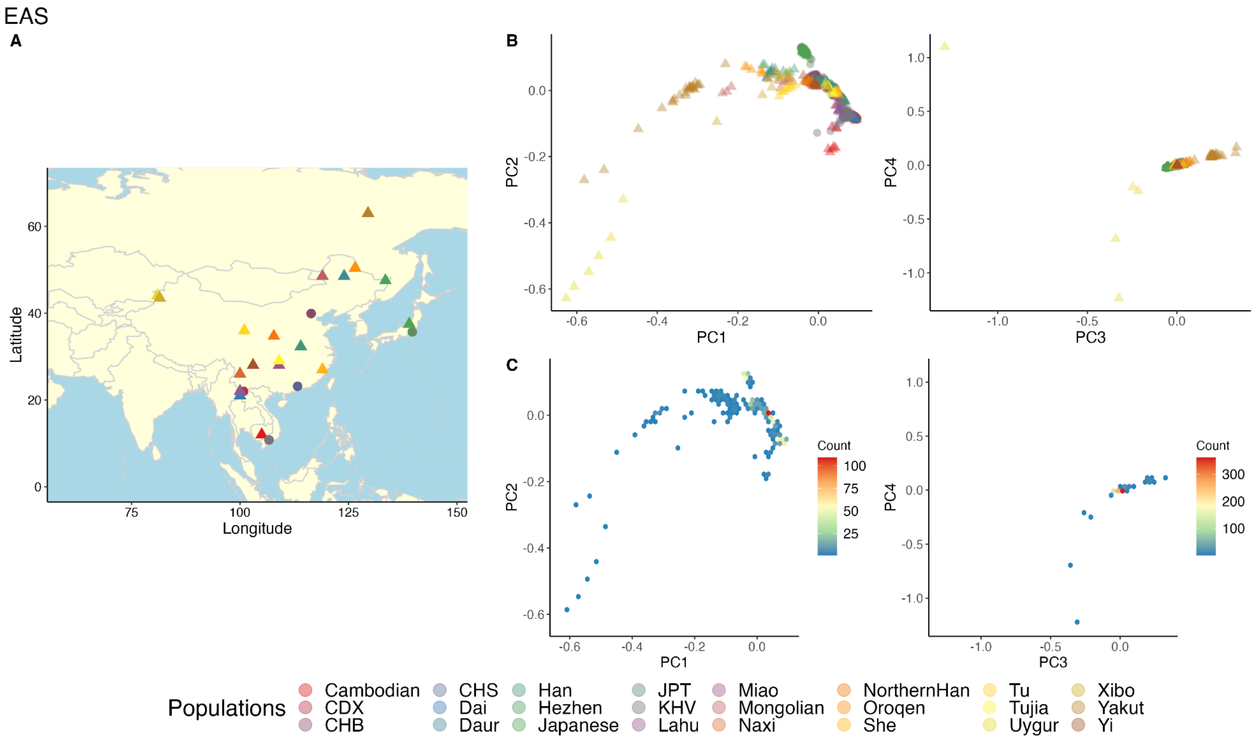


**Figure S12 | Subcontinental PCA in EAS populations.**

A) Map shows where all EAS samples in analyses are from. B) PCA biplots of PCs 1-4. PCA outliers were removed prior to this analysis. Filled circles indicate populations in the 1000 Genomes Project, while filled triangles indicate populations in HGDP. Population codes are as in **Table S1**. C) Density plot of PCA biplots of PCs 1-4.

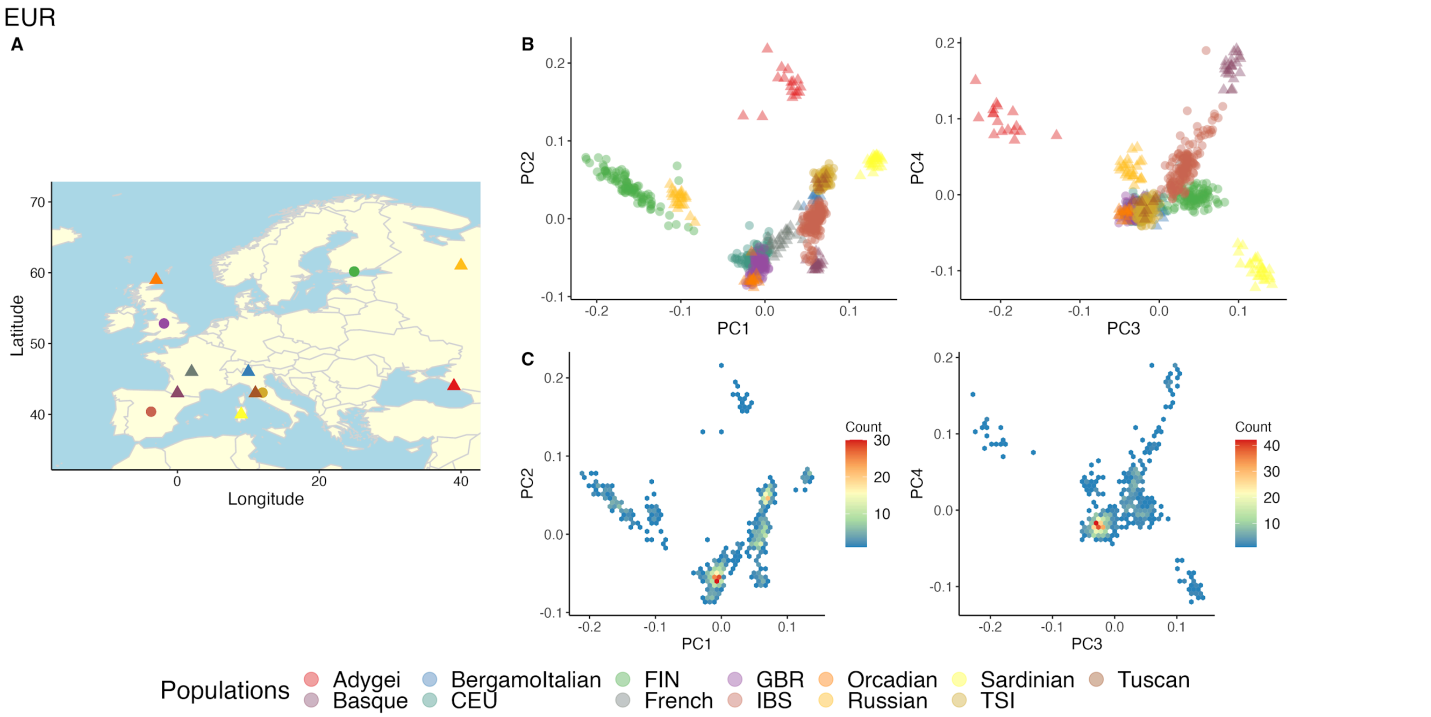


**Figure S13 | Subcontinental PCA in EUR populations.**
A) Map shows where all EUR samples in analyses are from. B) PCA biplots of PCs 1-4. PCA outliers were removed prior to this analysis. Filled circles indicate populations in the 1000 Genomes Project, while filled triangles indicate populations in HGDP. Population codes are as in **Table S1**. C) Density plot of PCA biplots of PCs 1-4.

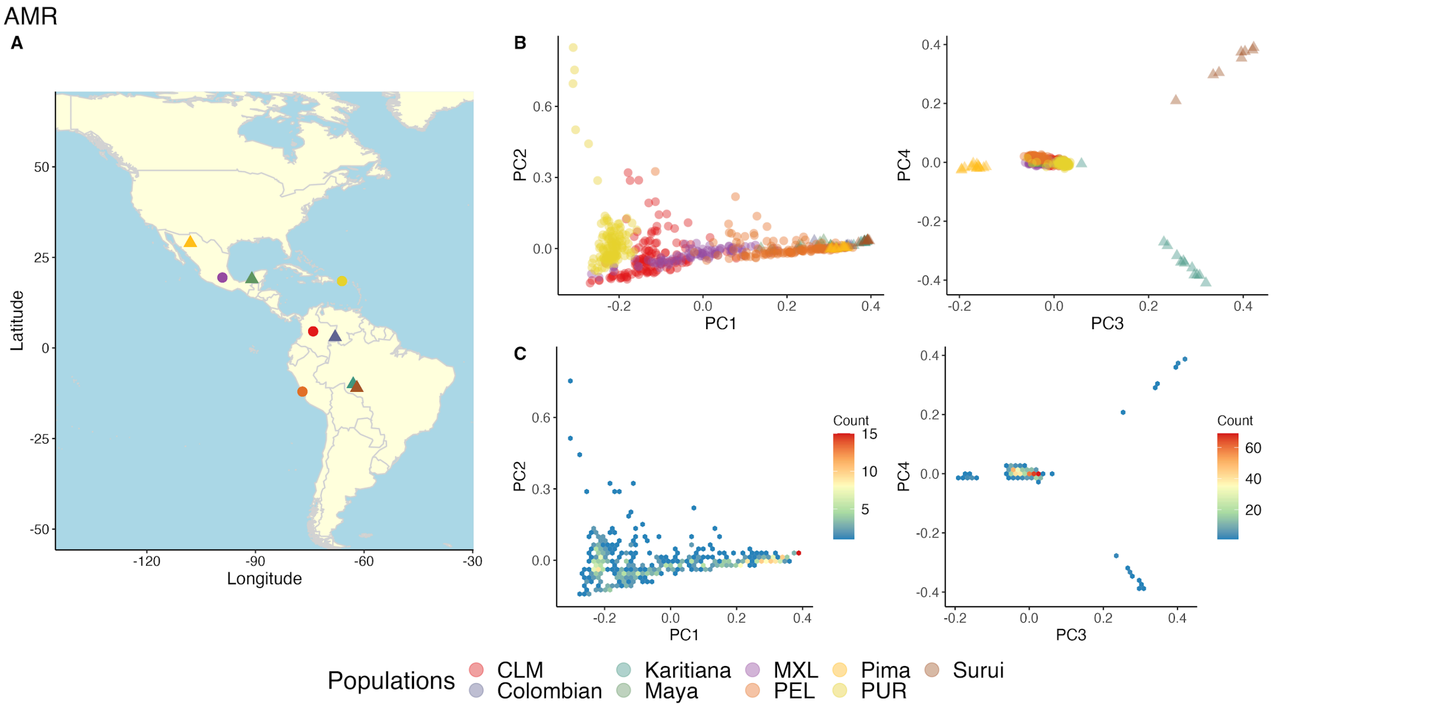


**Figure S14 | Subcontinental PCA in AMR populations.**
A) Map shows where all AMR samples in analyses are from. B) PCA biplots of PCs 1-4. PCA outliers were removed prior to this analysis. Filled circles indicate populations in the 1000 Genomes Project, while filled triangles indicate populations in HGDP. Population codes are as in **Table S1**. C) Density plot of PCA biplots of PCs 1-4.

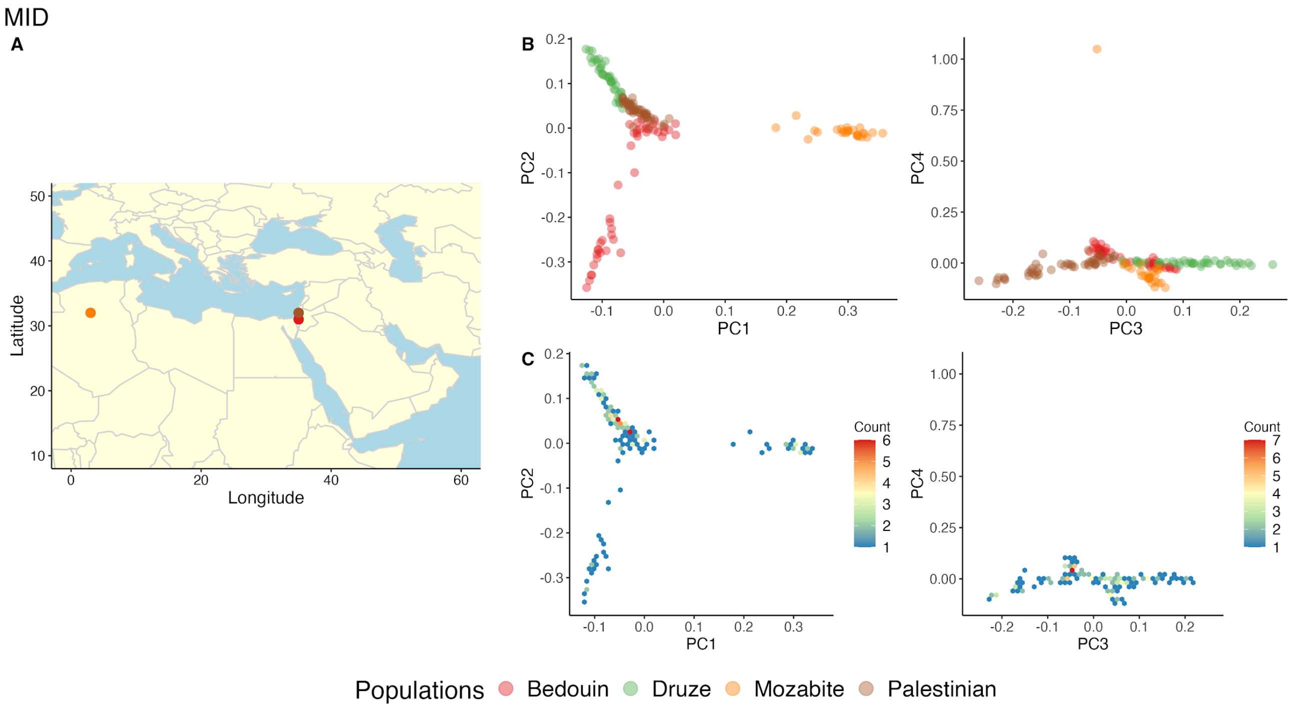


**Figure S15 | Subcontinental PCA in MID populations.**
A) Map shows where all MID samples in analyses are from. Palestinian and Druze have the same geographical coordinates. B) PCA biplots of PCs 1-4. PCA outliers were removed prior to this analysis. All MID populations are from HGDP. Population codes are as in **Table S1**. C) Density plot of PCA biplots of PCs 1-4.

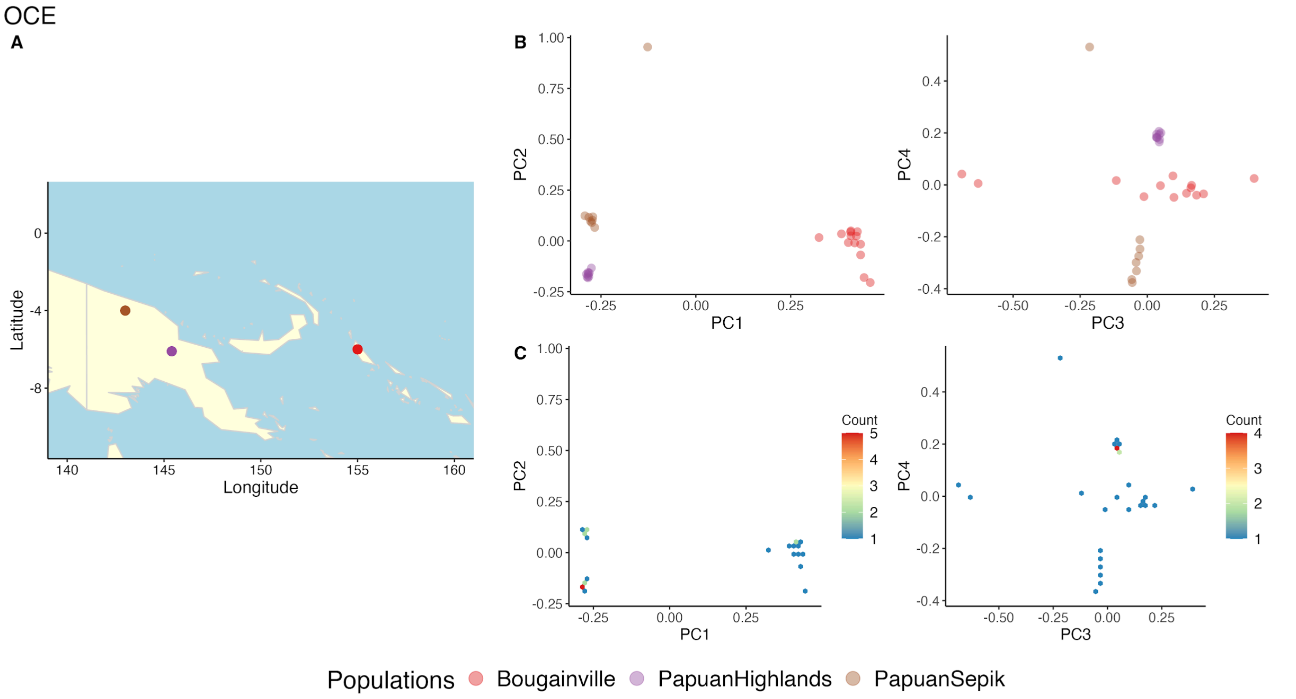


**Figure S16 | Subcontinental PCA in OCE populations.**
A) Map shows where all OCE samples in analyses are from. B) PCA biplots of PCs 1-4. PCA outliers were removed prior to this analysis. All OCE populations are from HGDP. Population codes are as in **Table S1**. C) Density plot of PCA biplots of PCs 1-4.

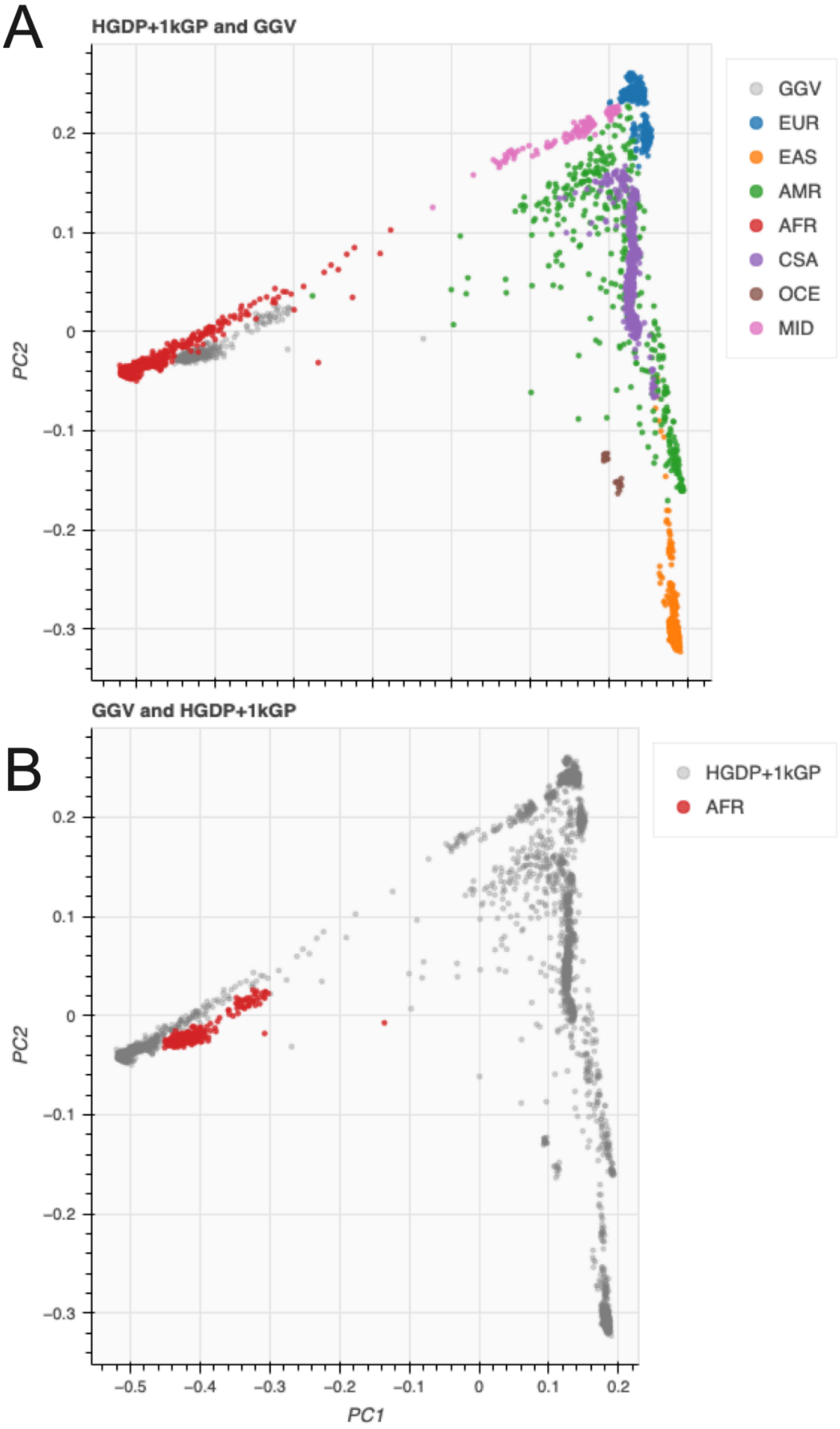


**Figure S17 | HGDP+1kGP ancestry labels applied to the Gambian Genome Variation (GGV) Project.** A) PCs 1 and 2 of all HGDP+1kGP samples with GGV projected into the same PC space, with each reference population colored and the GGV samples shown in grey. B) The same PCs with the reference data shown in grey and the GGV samples showing the assigned ancestry–all AFR.

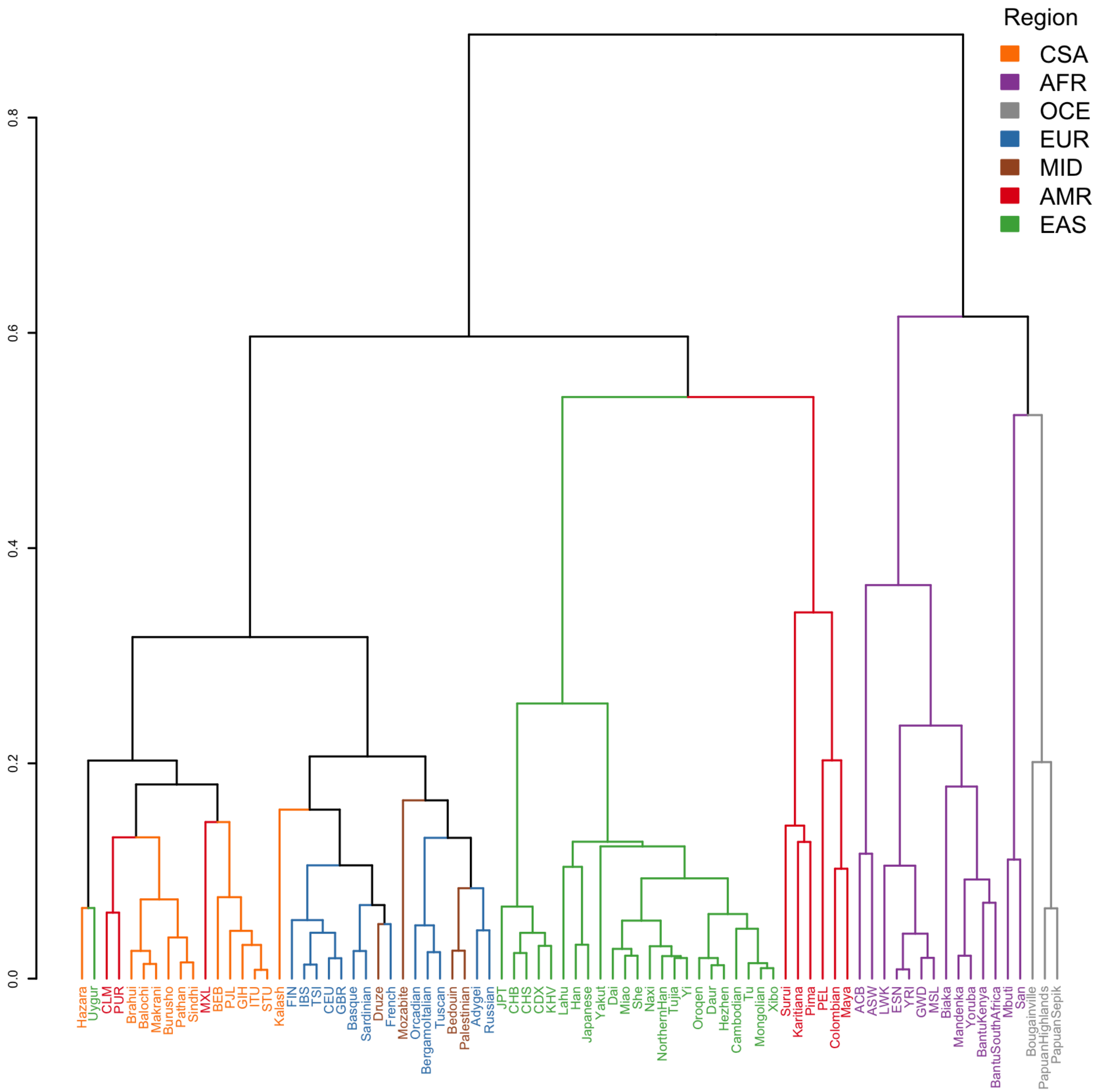


**Figure S18 | Dendrogram of the pairwise $F_{ST}$ heatmap between populations colored by geographical/genetic regions.**

Populations largely cluster by region with a few exceptions. MID and three AMR populations for example are interspersed among other regions.

**Table S7 | Populations interspersed among other geographical/genetic regions obtained from the pairwise $F_{ST}$ heatmap, colored by region.**

| Population | Region | Cluster |
|---|---|---|
| Hazara | CSA | Part of CSA population cluster |
| Uygur | EAS | EAS population among CSA cluster |
| CLM | AMR | 2 AMR populations among CSA cluster |
| PUR | AMR | |
| Brahui | CSA | CSA population cluster |
| Balochi | CSA | |
| Makrani | CSA | |
| Burusho | CSA | |
| Pathan | CSA | |
| Sindhi | CSA | |
| MXL | AMR | AMR population among CSA cluster |
| BEB | CSA | CSA population cluster |
| PJL | CSA | |
| GIH | CSA | |
| ITU | CSA | |
| STU | CSA | |
| Kalash | CSA | CSA population among EUR cluster |
| FIN | EUR | EUR population cluster |
| IBS | EUR | |
| TSI | EUR | |
| CEU | EUR | |
| GBR | EUR | |
| Basque | EUR | |
| Sardinian | EUR | |
| Druze | MID | MID population among EUR cluster |
| French | EUR | Part of EUR population cluster |
| Mozabite | MID | MID population among EUR cluster |
| Orcadian | EUR | EUR population cluster |
| BergamoItalian | EUR | |
| Tuscan | EUR | |
| Bedouin | MID | 2 MID populations among EUR cluster |
| Palestinian | MID | |
| Adygei | EUR | EUR population cluster |
| Russian | EUR | |
| JPT | EAS | EAS population cluster |
| CHB | EAS | |
| CHS | EAS | |
| CDX | EAS | |
| KHV | EAS | |
| Lahu | EAS | |
| Han | EAS | |
| Japanese | EAS | |
| Yakut | EAS | |
| Dai | EAS | |

| | | |
|---|---|---|
| Miao | EAS | |
| She | EAS | |
| Naxi | EAS | |
| NorthernHan | EAS | |
| Tujia | EAS | |
| Yi | EAS | |
| Oroqen | EAS | |
| Daur | EAS | |
| Hezhen | EAS | |
| Cambodian | EAS | |
| Tu | EAS | |
| Mongolian | EAS | |
| Xibo | EAS | |
| Surui | AMR | AMR population cluster |
| Karitiana | AMR | |
| Pima | AMR | |
| PEL | AMR | |
| Colombian | AMR | |
| Maya | AMR | |
| ACB | AFR | AFR population cluster |
| ASW | AFR | |
| LWK | AFR | |
| ESN | AFR | |
| YRI | AFR | |
| GWD | AFR | |
| MSL | AFR | |
| Biaka | AFR | |
| Mandenka | AFR | |
| Yoruba | AFR | |
| BantuKenya | AFR | |
| BantuSouthAfrica | AFR | |
| Mbuti | AFR | 2 AFR populations cluster with OCE |
| San | AFR | |
| Bougainville | OCE | OCE population cluster |
| PapuanHighlands | OCE | |
| PapuanSepik | OCE | |

**Table S8 | Pearson's correlation and Mantel tests results with and without waypoints.**
Waypoints and calculations are described further in Methods ("$F_{ST}$ versus geographical distance").

| Projects | With waypoints | | Without waypoints | |
|---|---|---|---|---|
| | Pearson's correlation coefficient | Mantel statistic | Pearson's correlation coefficient | Mantel statistic |
| HGDP | 0.7615882 | 0.5471567 | 0.7095223 | 0.4504546 |
| 1kGP | 0.3672138 | 0.1168814 | 0.327703 | 0.1009129 |
| Cross-project | 0.450008 | 0.1925107 | 0.4119236 | 0.1549286 |
| Everything | 0.5606575 | 0.3084334 | 0.5290618 | 0.2592783 |

#### Quality control

Our sample QC procedure was mostly the same as in gnomAD, but differed slightly. Specifically, because whole populations were removed from gnomad ‘fail_’ filters, we did not filter on the basis of these, which were used in gnomAD v3.1. The clearest example of filters that failed was the fail_n_snp_residual filter, as shown in **Figure S19**.

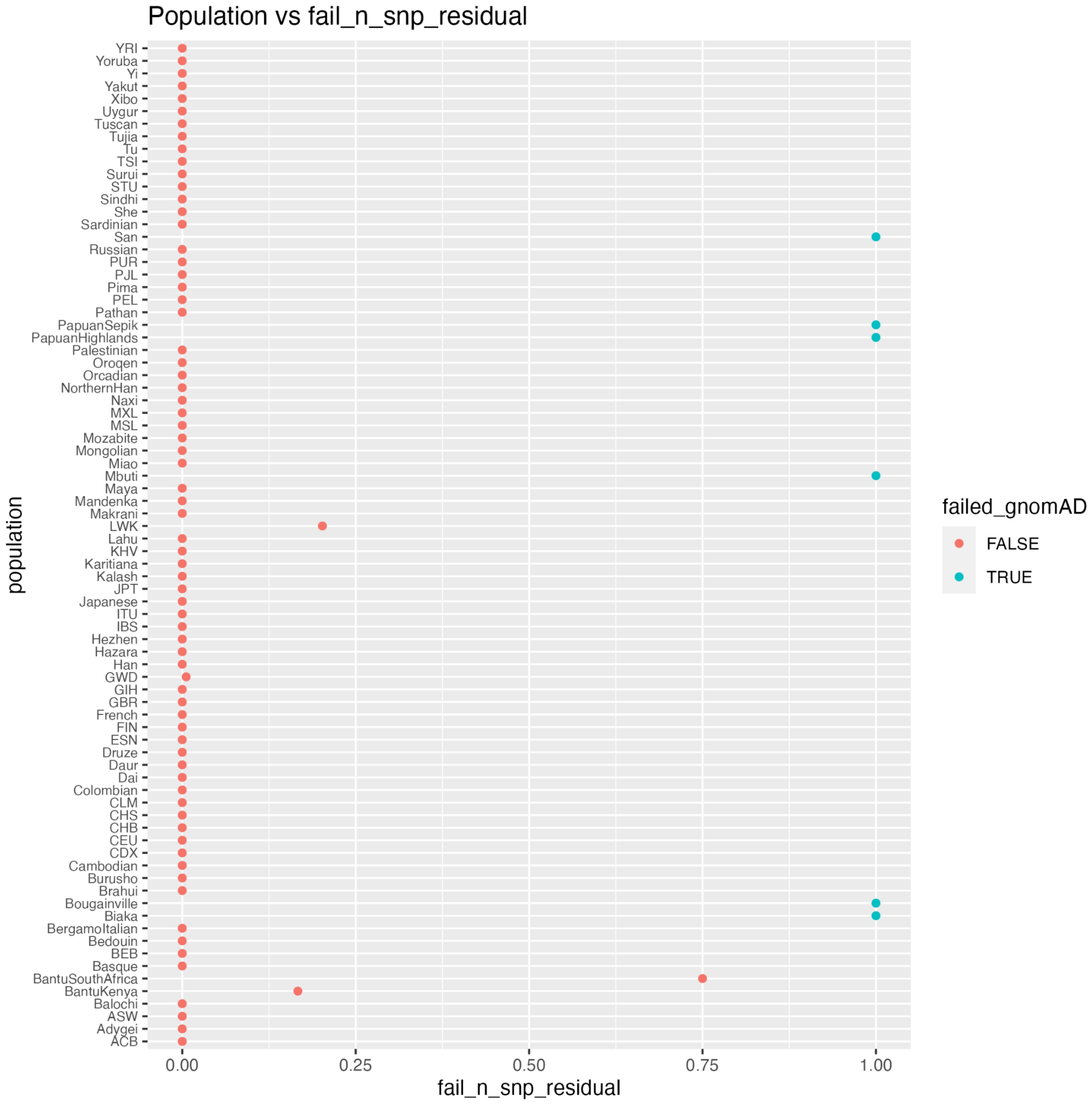


**Figure S19 | Example of a filter that was included in gnomAD v3.1 but excluded from this project.** The “fail_n_snp_residual” filter, which regresses out principal components from the number of SNPs in an effort to identify technical outliers, would have excluded whole continental groups and populations in this resource because these groups are distinct from the majority of individuals in gnomAD.

**Table S9 | gnomAD Sample and Variant Quality Control Steps**

This table contains the steps conducted by the gnomAD team on the jointly called HGDP+1kGP dataset. These QC steps were conducted prior to the additional quality control steps we conducted as outlined in the methods section. Samples and variants that passed gnomAD's QC were labeled as PASS. For additional details on the QC steps conducted, see the gnomAD blogpost on the release of the HGDP+1kGP dataset (https://gnomad.broadinstitute.org/news/2020-10-gnomad-v3-1-new-content-methods-annotations-and-data-availability/#sample-and-variant-quality-control).

**Sample QC Steps**

Hard Filtering
Using sample QC metrics computed using Hail's sample_qc() module on all autosomal bi-allelic SNVs the following filters were applied:

- Number of SNVs: < 2.4M or > 3.75M
- Number of singletons: >100k
- Ratio of heterozygous to homozygous variants: > 3.3

Hard filtering using BAM-level metrics was performed when such metrics were available. Removed samples with:

- Contamination: > 5%
- Chimeras: > 5%
- Median insert size: < 250

---

Sex Inference
Used a rough F-stat cutoff of 0.5 to split samples into XX and XY categories. Final X and Y ploidy cutoffs were determined from the means and standard deviations of those XX and XY distributions. Sex was assigned based on the following cutoffs:

- XY:
    - Normalized X coverage < 1.29 &
    - Normalized Y coverage > 0.1 &
    - Normalized Y coverage < 1.16
- XX:
    - Normalized X coverage > 1.45 &
    - Normalized X coverage < 2.4 &
    - Normalized Y coverage < 0.1

---

Ancestry inference
30 principal components (PCs) were computed using principal components analysis (PCA) using Hail's hwer_normalized_pca() function. 76,419 high quality variants were selected for these analyses using the following selection criteria:

1. Lifting over all sites from gnomAD v2.1 to GRCh38
2. Add ~5k sites widely used for quality control of GWAS data defined by Shaun Purcell and lifted them over to GRCh38
3. From the two above sets, selected all bi-allelic SNVs with an Inbreeding coefficient > -0.25 (no excess of heterozygotes)

The 30 PCs were visually inspected to determine which PCs were capturing variants explained by the available "known" population labels. The first 16 PCs were chosen as best capturing the global ancestry variation and then a random forest classifier was trained using those PCs as features on the samples with "known" population labels. Ancestry labels were then assigned to all samples for which the random forests model, and all remaining samples were given a population label of "other" (oth)

---

Sample QC metric outlier filtering
Computed sample QC metrics using Hail's sample_q() module and regressed out the first 8 ancestry assignment PCs. Samples which fell outside the 4 median absolute deviations (MADs) from the median for the following metrics:

- Number of snps
- Transition/transversion ratio
- Insertion/deletion ratio

- Number of insertions
- Number of deletions
- Number of heterozygotes
- Number of homozygous variants
- Number of transitions
- Number of transversions

Additionally filtered samples over 8 MADs above the median number of singletons and over 4 MADs above the median ratio of heterozygous/homozygous variants call ratio.

**Variant QC**

Performed variant QC using the allele-specific version of GATK Variant Quality Score Recalibration (VQSR) with the following features:

- SNVs: AS_FS, AS_SOR, AS_ReadPosRankSum, AS_MQRankSum, AS_QD, AS_MQ
- Indels: AS_FS, AS_SOR, AS_ReadPosRankSum, AS_MQRankSum, AS_QD

The standard GATK training resources (HapMap, Omni, 1000 Genomes, Mills indels) were used in addition to ~19M transmitted singletons (alleles confidently observed exactly twice in gnomAD, once in a parent and once in a child) from 6,743 trios present in the raw data.

<u>Variant hard filters</u>

- No high-quality genotype (GQ >=20, DP>=10, and AB>=0.2 for heterozygotes) called for the variant
- Inbreeding Coefficient < -0.3

#### Dataset Comparisons

**Table S10 | Filters Applied to Comparison Datasets**
For each dataset, the filter name is written along with the number of variants removed by that filter. Phase3 1kGP is not included in this table as there were no filters listed for that dataset.

| Dataset | Filter Name | Number of Variants Removed | Number of Variants Remaining after Filters and Removing Chromosomes X and Y |
|---|---|---|---|
| gnomAD v3 | AC0 | 32,857,346 | 644,267,978 |
| | AC0, AS_VQSR | 44,883,875 | |
| | AC0, AS_VQSR, InbreedingCoeff | 471 | |
| | AC0, InbreedingCoeff | 457 | |
| | AS_VQSR | 37,019,424 | |
| | AS_VQSR, Inbreeding Coeff | 105,015 | |
| | InbreedingCoeff | 167,701 | |
| Bergstrom HGDP | ExcHet | 211,310 | 75,708,475 |
| | ExcHet, LOW_VQSLOD | 253,580 | |
| | LOW_VQSLOD | 3,813,640 | |
| Phase 3 1kGP | N/A | 0 | 73,159,510 |
| NYGC 1kGP | VQSRTrancheINDEL99.00to100.00 | 4,118,436 | 116,715,566 |
| | VQSRTrancheSNP99.80to100.00 | 6,790,855 | |

**Table S11 | Variants absent in HGDP+1kGP that were identified in comparison datasets depending on whether variants passed QC in each dataset.**

The numbers in this table are the counts of variants which are only in the comparison datasets, listed in the first column. Counts were generated by taking the disjoint subset of the pre- and post-QC HGDP+1kGP dataset with both the pre- and post-QC versions of the comparison dataset. For the comparison datasets, QC is defined as applying the filters supplied by the datasets as in **Table S10** as well as removing the X and Y chromosome from the dataset. For HGDP+1kGP, QC is defined as applying gnomAD sample, variant, and genotype QC filters, as well as removing PCA outliers. For the gnomAD dataset only, the post-QC version of HGDP+1kGP includes PCA outliers since gnomAD did not filter these ancestry outliers. Note that for Phase 3 1kGP, since there were no QC filters to apply, the pre- and post-QC numbers only refer to pre and post removal of the X and Y chromosomes.

| | | | HGDP+1kGP | |
|---|---|---|---|---|
| | | | Pre-QC | Post-QC |
| gnomAD v3* | Pre-QC | SNVs | 499,396,055 | 521,411,946 |
| | | Indels | 78,053,396 | 88,302,017 |
| | | Total | 577,449,451 | 609,713,963 |
| | Post-QC | SNVs | 518,873,266 | 422,293,234 |
| | | Indels | 85,396,533 | 47,665,075 |
| | | Total | 604,269,799 | 469,958,309 |
| Bergstrom HGDP | Pre-QC | SNVs | 2,422,849 | 6,790,567 |
| | | Indels | 258,675 | 1,936,519 |
| | | Total | 2,681,524 | 8,727,086 |
| | Post-QC | SNVs | 1,910,423 | 4,928,546 |
| | | Indels | 89,153 | 741,002 |
| | | Total | 1,999,576 | 5,669,548 |
| Phase 3 1kGP | Pre-QC | SNVs | 1,555,765 | 2,337,154 |
| | | Indels | NA | NA |
| | | Total | 1,555,765 | 2,337,154 |
| | Post-QC | SNVs | 1,552,222 | 2,330,829 |
| | | Indels | NA | NA |
| | | Total | 1,552,222 | 2,330,829 |
| NYGC 1kGP | Pre-QC | SNVs | 4,538,405 | 11,241,168 |
| | | Indels | 265,914 | 2,249,456 |
| | | Total | 4,804,319 | 13,490,624 |
| | Post-QC | SNVs | 1,449,073 | 5,369,909 |
| | | Indels | 155,352 | 1,303,555 |
| | | Total | 1,604,425 | 6,673,464 |

**Table S12 | Variant counts and percentages of comparison datasets.**
The number of variants in are calculated for each minor allele frequency (MAF) bin. The MAF is represented as the MAF in the HGDP+1kGPd dataset. Variants with a MAF of 0% are not in the HGDP+1kGP dataset. For each comparison dataset there are counts for the number of variants in the comparison dataset only, in both the comparison dataset and HGDP+1kGP, and in HGDP+1kGP only. For each dataset at the bottom of each total count is the percentage of variants in each category previously described.

| MAF in HGDP+1kGP | Variants in Comparison Only | Variants in Both | Variants in HGDP+1kGP Only |
|---|---|---|---|
| gnomAD v3 | | | |
| 0% | 469,958,679 | - | - |
| 0.01%-0.1% | - | 74,959,954 | 37,645,877 |
| 0.1%-1% | - | 25,783,919 | 1,693,102 |
| 1.0%-10% | - | 11,708,163 | 11,340 |
| 10-50% | - | 7,533,718 | 3,074 |
| **Total** | 469,958,679 (75%) | 119,985,754 (19%) | 39,353,393 (6%) |
| Bergstrom HGDP | | | |
| 0% | 5,669,548 | - | - |
| 0.01%-0.1% | - | 37,191,483 | 77,124,525 |
| 0.1%-1% | - | 16,775,062 | 5,203,182 |
| 1.0%-10% | - | 9,304,753 | 1,063,657 |
| 10-50% | - | 6,767,625 | 464,564 |
| **Total** | 5,669,548 (4%) | 70038,923 (44%) | 83,855,928 (53%) |
| Phase3 1kGP | | | |
| 0% | 2,330,829 | - | - |
| 0.01%-0.1% | - | 44,952,062 | 69,363,946 |
| 0.1%-1% | - | 14,177,436 | 7,800,808 |
| 1.0%-10% | - | 6,599,319 | 3,769,091 |
| 10-50% | - | 5,099,864 | 2,132,325 |
| **Total** | 2,330,829 (1%) | 70,828,681 (45%) | 83,066,170 (53%) |
| NYGC 1kGP | | | |

| | | | |
|---|---|---|---|
| 0% | 6,673,464 | - | - |
| 0.01%-0.1% | - | 75,666,785 | 38,649,223 |
| 0.1%-1% | - | 18,481,913 | 3,496,331 |
| 1.0%-10% | - | 9,201,326 | 1,167,084 |
| 10-50% | - | 6,692,078 | 540,111 |
| **Total** | 6,673,464<br>(4%) | 110,042,102<br>(69%) | 43,852,749<br>(27%) |

#### Analysis tutorials

To show examples of how to use the individual-level data in a cloud-computing environment, we have created a series of tutorials in iPython notebooks that make use of Hail. These tutorials show how to merge datasets, apply sample and variant QC, run ancestry analysis via PCA and visualization, generate summary statistics of genomes by population, compute and plot population divergence statistics via $F_{ST}$ and $F_2$ statistics, and intersect external datasets with this dataset and infer ancestry information using project meta-data. The organization of these notebooks is outlined in **Figure 6**.

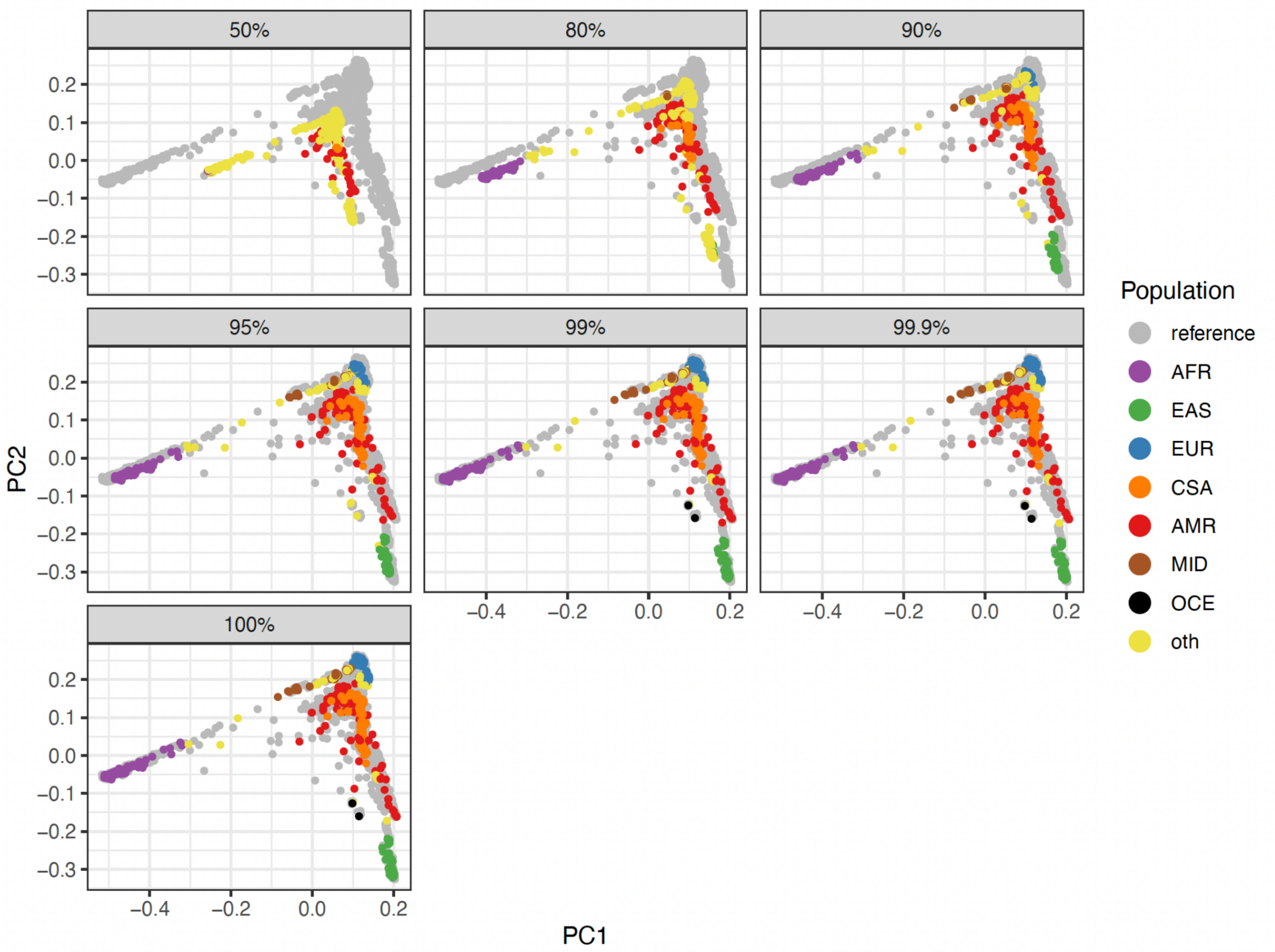


**Figure S20 | PCA shrinkage analysis to determine acceptable levels of missingness before ancestry resolution becomes too low to accurately assign population labels.**
We started with a set of SNPs that were used in other PCA (e.g. **Figure 2**), which had undergone minor allele frequency filtering, missingness filtering, and LD pruning. We randomly selected 80% of samples (N=2,720) to train the random forest with corresponding meta-data labels as usual and held out 20% of samples as a test dataset (N=680). After filtering out monomorphic sites from the training dataset once samples were divided, we retained 200,403 variants which were used to train the random forest. We randomly downsampled SNPs in the test dataset to include 50%, 80%, 90%, 95%, 99%, 99.9%, and 100% of SNPs in the training dataset. These plots show the corresponding projected PCs in the test dataset, showing the extent to which shrinkage affects analyses. **Table S13** shows rates of unclassified individuals by SNP missingness in the test dataset.

**Table S13 | Shrinkage analysis matches and no classification numbers by SNP missingness in the test dataset, as shown in Figure S20.**
There were no mismatched labels assigned.

| Fraction of SNPs in test dataset out of training dataset | Match | No assignment |
|---|---|---|
| 1 | 653 / 680 = 0.96 | 27 / 680 = 0.04 |
| 0.999 | 654 / 680 = 0.96 | 26 / 680 = 0.04 |
| 0.99 | 650 / 680 = 0.96 | 30 / 680 = 0.04 |
| 0.95 | 624 / 680 = 0.92 | 56 / 680 = 0.08 |
| 0.9 | 563 / 680 = 0.83 | 117 / 680 = 0.17 |
| 0.8 | 340 / 680 = 0.5 | 340 / 680 = 0.5 |
| 0.5 | 102 / 680 = 0.15 | 578 / 680 = 0.85 |
